# MSL3 coordinates a transcriptional and translational meiotic program in female Drosophila

**DOI:** 10.1101/2019.12.18.879874

**Authors:** Alicia McCarthy, Kahini Sarkar, Elliot T Martin, Maitreyi Upadhyay, Joshua R James, Jennifer M Lin, Seoyeon Jang, Nathan D Williams, Paolo E Forni, Michael Buszczak, Prashanth Rangan

**Author notes:** Corresponding Author and Lead Contact: Prashanth Rangan, Department of Biological Sciences/RNA Institute, University at Albany, SUNY, 1400 Washington Avenue, Albany, NY 12222.

## Abstract

Gamete formation from germline stem cells (GSCs) is essential for sexual reproduction. However, the regulation of GSC differentiation and meiotic entry are incompletely understood. Set2, which deposits H3K36me3 modifications, is required for differentiation of GSCs during *Drosophila* oogenesis. We discovered that the H3K36me3 reader Male-specific lethal 3 (MSL3) and the histone acetyltransferase complex Ada2a-containing (ATAC) cooperate with Set2 to regulate entry into meiosis in female *Drosophila*. MSL3 expression is restricted to the mitotic and early meiotic stages of the female germline, where it promotes transcription of genes encoding synaptonemal complex components and a germline enriched *ribosomal protein S19* paralog, *RpS19b*. *RpS19b* upregulation is required for translation of Rbfox1, a known meiotic cell cycle entry factor. Thus, MSL3 is a master regulator of meiosis, coordinating the expression of factors required for recombination and GSC differentiation. We find that MSL3 is expressed during mouse spermatogenesis, suggesting a conserved function during meiosis.

## Introduction

Germ cells give rise to gametes, a fundamental requirement for sexual reproduction. The production of gametes is tightly controlled to ensure a constant supply throughout the reproductive life of an organism (Cinalli et al., 2008; Kimble, 2011; Lehmann, 2012; Spradling et al., 1997). Germ cells can directly differentiate to enter meiosis or become germline stem cells (GSCs) (Edson et al., 2009; Fuller and Spradling, 2007; Nikolic et al., 2016; Saitou and Yamaji, 2010; Sharma et al., 2019). GSCs divide mitotically to both self-renew and generate differentiating daughters that can enter meiosis (Fayomi and Orwig, 2018; Fuller and Spradling, 2007; Kimble, 2011; Lehmann, 2012; De Rooij, 2017; Spradling et al., 2011). Loss of meiotic entry results in infertility (Cohen et al., 2006; Handel and Schimenti, 2010; Hughes et al., 2018; Lesch and Page, 2012; Marston and Amon, 2004; Soh et al., 2015), so propagation of sexually reproducing organisms hinges upon the ability of the germ cells to enter meiosis.

In mammals, the entry into meiosis is promoted by steroid signaling during oogenesis and spermatogenesis (Bowles and Koopman, 2007; Griswold et al., 2012). Retinoic acid (RA) from somatic cells activates the transcription factor *Stimulated by retinoic acid 8* (*Stra8*) in the germline (Bowles et al., 2006; Endo et al., 2015, 2017; Koubova et al., 2006; Oulad-Abdelghani et al., 1996; Zhou et al., 2008). During spermatogenesis, STRA8 fosters meiotic entry by promoting transcription of a broad gene expression program (Abby et al., 2016; Bailey et al., 2017; Jain et al., 2018; Kojima et al., 2019; Soh et al., 2017). However, *Stra8* is not sufficient to induce meiosis, suggesting a cell type-specific chromatin landscape and/or factors that cooperate with STRA8 (Endo et al., 2015; Kojima et al., 2019; Miyauchi et al., 2017; Zhou et al., 2008). In addition, *Stra8* is not conserved outside of vertebrates (Fujiwara and Kawamura, 2003; Hickford et al., 2017). The transcriptional machinery that promotes meiotic entry in other organisms has remained elusive.

Meiotic differentiation is well characterized in *Drosophila* (Hales et al., 2015). In both male and female *Drosophila*, germ cells acquire a GSC fate prior to differentiating into gametes (Dansereau and Lasko, 2008; Lehmann, 2012; Marlow, 2015). *Drosophila* ovaries are composed of individual egg producing units called ovarioles. A structure called the germarium lies at the tip of each ovariole and houses 2-3 GSCs, which are marked by round organelles called spectrosomes (Eliazer and Buszczak, 2011; Kahney et al., 2019; Kirilly et al., 2011; Morris and Spradling, 2011; Morrison and Spradling, 2008; Spradling et al., 2001, 2011, 2008; Xie, 2000; Xie and Spradling, 2000) (Figure 1A). GSCs both self-renew and differentiate into cystoblasts (CBs) that divide without cytokinesis to give rise to 2-, 4-, 8-, and 16-cell cysts, which are marked by branched structures called fusomes (Chen and McKearin, 2003a, 2003b; Xie, 2013).

**Figure 1.**
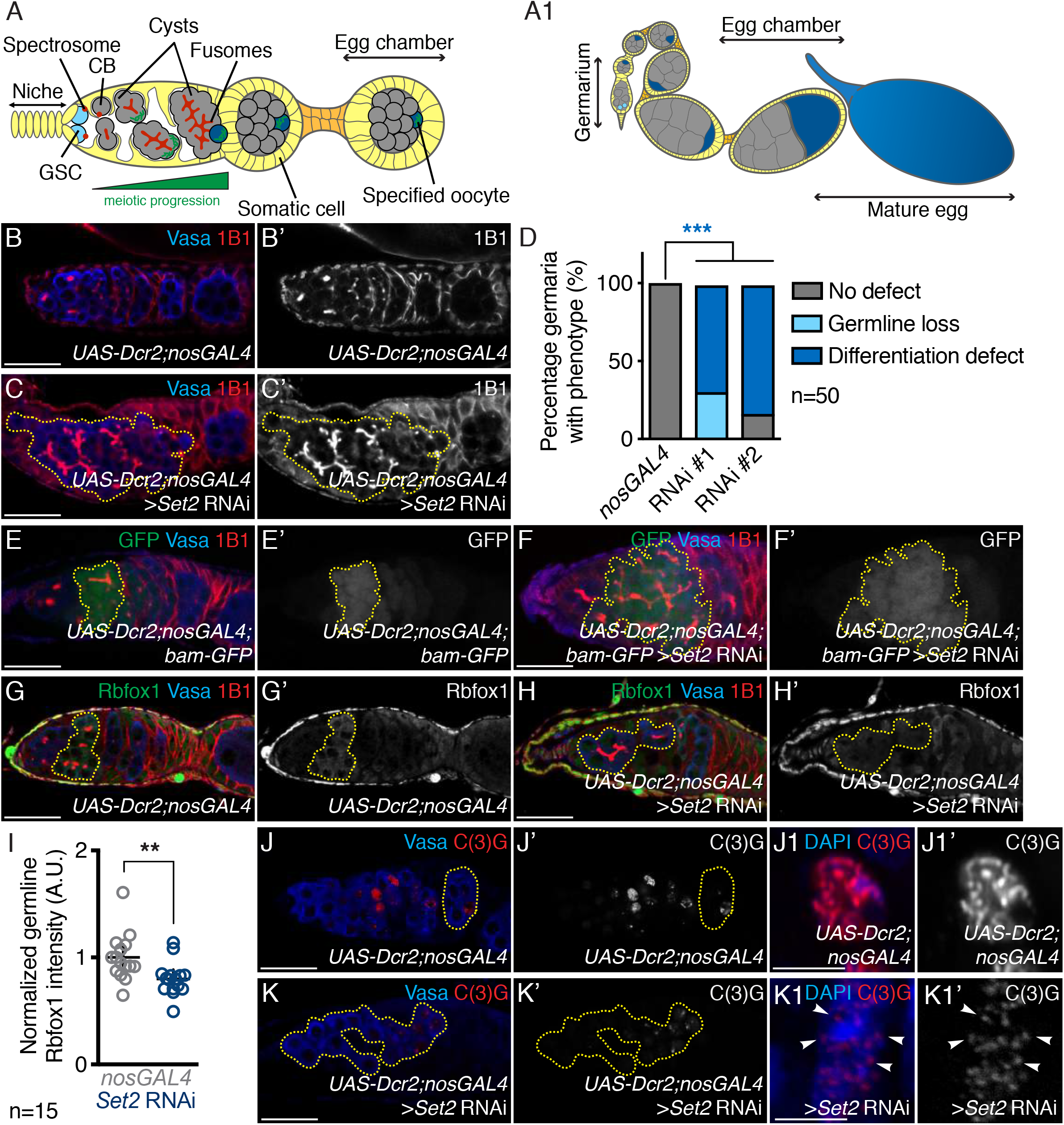
Set2 is required in the germline for meiotic progression during oogenesis. (A) A schematic of a *Drosophila* germarium where germ cells (gray, light and dark blue) are surrounded by somatic cells (yellow). The germline stem cells (GSCs; light blue) reside near a somatic niche (yellow). The GSC divides to give rise to daughter cells called cystoblasts (CBs; gray). Both GSCs and CBs are marked by round structures called spectrosomes (red). CBs turn on a differentiation program and will undergo incomplete mitotic divisions, giving rise to 2, 4, 8, and 16-cell cysts (gray), marked by branched structures called fusomes (red). During the cyst stages germ cells progress through meiotic prophase I (green rectangles, green triangle below). Upon 16-cell cyst formation, a single cell will be specified as the oocyte (dark blue) while the other 15 cells become support cells called nurse cells (gray). The 16-cell cyst will migrate, bud off from the germarium, be encapsulated by the soma (yellow), and generate egg chambers. (A1) A schematic of a *Drosophila* ovariole. The ovariole consists of egg chambers that are discrete stages of development, connected by somatic cells (orange). As egg chambers develop, they increase in size and house the maturing oocyte eventually giving rise to a mature egg (blue). (B-B’) Control and (C-C’) germline depleted *Set2* (RNAi line #1) germaria stained for Vasa (blue) and 1B1 (red) shows that *Set2* germline depletion results in irregular cysts (70% in *Set2* RNAi line #1 and 84% in *Set2* RNAi line #2 compared to 0% in *nosGAL4*; *p=4.1E-15* and *p*<*2.2E-16*, respectively, n=50) (yellow dashed outline) and germline loss (30% in *Set2* RNAi line #1 and 0% in *Set2* RNAi line #2 compared to 0% in *nosGAL4*; *p*=*3.2E-12* and *p=1*, respectively, n=50). 1B1 channel is shown in B’ and C’. Quantitation in (D), statistical analysis performed with Fisher’s exact test on differentiation defect; *** indicates *p<0.001*. (E-E’) Control and (F-F’) germline depleted *Set2* germaria both carrying a *bam-GFP* transgene stained for GFP (green), Vasa (blue), and 1B1 (red) shows that *Set2* germline depletion results in irregular GFP positive cysts compared to control (yellow dashed outline) (90% in *Set2* RNAi compared to 4% in *nosGAL4*; *p*<*2.2E-16*, n=50). Statistical analysis performed with Fisher’s exact test. GFP channel is shown in E’ and F’. (G-G’) Control and (H-H’) germline depleted *Set2* germaria stained for Rbfox1 (green), Vasa (blue), and 1B1 (red) shows that *Set2* germline depletion results in decreased levels of Rbfox1 in the germline compared to control (yellow dashed outline) (0.7±0.1 in *Set2* RNAi compared to 1.0±0.1 in *nosGAL4*; *p=0.0076*, n=15). Rbfox1 channel is shown in G’ and H’. Quantitation in (I), statistical analysis performed with Student t-test; ** indicates *p<0.01*. (J-J’ and J1-J1’) Control and (K-K’ and K1-K1’) germline depleted *Set2* germaria stained for Vasa (blue) and C(3)G (red) shows that *Set2* germline depletion results in aberrant C(3)G staining compared to control (yellow dashed outline) (100% in *Set2* RNAi compared to 2% in *nosGAL4*; *p<2.2E-16*, n=50) and improper assembly of the synaptonemal complex (white arrows). Statistical analysis performed with Fisher’s exact test. C(3)G channel is shown in J’, J1’, K1, and K1’. Scale bar for J1-J1’ and K1-K1’ is 2 μm, scale bar for all other images is 20 μm.

The somatic niche of the germarium provides Decapentaplegic (DPP) signaling that leads to phosphorylation of Mothers against DPP (pMad) in GSCs, and transcriptional repression of the differentiation factor *bag of marbles* (*bam*) (Chen and McKearin, 2003a, 2003b; Kai and Spradling, 2003). After GSC division, the CB is displaced from the niche, allowing for Bam expression (Chen and McKearin, 2003a, 2003b). Bam is sufficient to promote the transition from CB to a differentiated 8-cell cyst (McKearin and Ohlstein, 1995; McKearin and Spradling, 1990).

In the 8-cell cyst, expression of the cytoplasmic isoforms of RNA-binding Fox protein 1 (Rbfox1) leads to translational downregulation of self-renewal factors to promote expression of Bruno (Bru) (Carreira-Rosario et al., 2016; Tastan et al., 2010). Bru, in turn, translationally represses mitotic factors, which promote cyst divisions, and regulates entry into a meiotic cell cycle (Parisi et al., 2001; Sugimura and Lilly, 2006; Wang and Lin, 2007). Multiple cells in the cysts initiate meiosis, but only the oocyte will commit to meiosis; the other 15 cells acquire a nurse cell fate in the 16-cell cyst stage (Carpenter, 1975, 1994; Carpenter and Sandler, 1974; Huynh and St Johnston, 2004; Mach and Lehmann, 1997; Navarro et al., 2001; Theurkauf et al., 1993). The oocyte and the 15 nurse cells are encapsulated by somatic cells to form a developing egg chamber and eventually an egg (Figure 1A1). Although Rbfox1 expression in the germline is essential for entry into a meiotic cell cycle and oocyte specification, how it is induced is unclear (Carreira-Rosario et al., 2016).

Another hallmark of meiosis, apart from a specialized cell cycle, is homologous chromosome recombination. This process is regulated by the formation of the synaptonemal complex (SC) (Ables, 2015; Carpenter, 1975; Hughes et al., 2018). The SC starts to assemble on homologs in up to four nuclei (Takeo et al., 2011), but is maintained only in the specified oocyte (Page and Hawley, 2001; Von Stetina and Orr-Weaver, 2011) (Figure 1A). How transcription of SC components is activated during meiotic commitment is not well understood.

GSC differentiation during *Drosophila* oogenesis requires the histone methyltransferase SET domain containing 2 (Set2), which confers histone H3 lysine 36 trimethylation (H3K36me3) (Larschan et al., 2007; Mukai et al., 2015). H3K36me3 typically marks transcriptionally active genes (Bannister and Kouzarides, 2011; Dong and Weng, 2013; Keogh et al., 2005). How H3K36me3 regulates GSC differentiation is not clear. Interestingly, in male *Drosophila*, H3K36me3 facilitates recognition of the X chromosome by the Male-Specific Lethal (MSL) complex, which leads to hyper-transcription of the male X and gene dosage compensation with females, which have two X chromosomes (Bell et al., 2008; Conrad et al., 2012a; Larschan et al., 2007; Samata and Akhtar, 2018; Sural et al., 2008). Within the MSL complex, the chromodomain (CD) of MSL3 reads the H3K36me3 marks and the histone acetyl transferase (HAT) Males absent on the first (MOF) drives acetylation of histone H4 lysine 16 (H4K16ac) (Bone et al., 1994; Conrad et al., 2012b; Gu et al., 2000; Hilfiker et al., 1997; Kadlec et al., 2011; Larschan et al., 2007; Sural et al., 2008; Turner et al., 1992). Female flies do not assemble the MSL complex because some key components are not expressed (Bachiller and Sánchez, 1989; Bashaw and Baker, 1997; Belote, 1983; Belote and Lucchesi, 1980; Kelley et al., 1997; Uchida et al., 1981). MSL proteins are conserved in mammals and regulate embryonic stem cell differentiation (Basilicata et al., 2018; Chelmicki et al., 2014; Heard and Disteche, 2006; Keller and Akhtar, 2015; Laverty et al., 2010; Ravens et al., 2014). However, if MSL proteins in *Drosophila* regulate gene expression beyond their role in dosage compensation is not known.

Here, we identified a transcriptional axis that regulates the transition to the meiotic program in *Drosophila*. We find that Set2, MSL3, and a HAT complex, Ada2a containing (ATAC), mediate progression into meiosis. We discovered that MSL3 is expressed in the pre-meiotic and early meiotic stages during oogenesis, where it promotes the HAT-mediated transcription of several members of the SC as well as a germline-specific paralog of eukaryotic *Ribosomal protein S19* (*eRpS19*/*RpS19*). In humans, mutations of RpS19 disrupt hematopoiesis due to translational dysregulation of distinct mRNAs (Draptchinskaia et al., 1999; Ludwig et al., 2014; Willig et al., 2000). We discovered that expression of RpS19b helps increase overall levels of RpS19, which is then required for translation of *Rbfox1* and thus entry into meiosis in female flies. We show that MSL3 is also expressed in differentiating mouse spermatogonia, downstream of the critical meiosis promoting factor STRA8, suggesting an evolutionarily conserved role for MSL3 in promoting meiosis. Thus, the Set2-MSL3-ATAC axis directly regulates transcription and indirectly regulates translation of key meiotic factors to promote proper GSC differentiation.

## Results

### Set2 is required in the germline for meiotic progression during oogenesis

To determine how Set2 promotes oogenesis in *Drosophila*, we stained control and *Set2* depleted fly gonads with antibodies against Vasa, a germline marker, and 1B1, a marker of somatic cell membranes, spectrosomes, and fusomes. Compared to controls, *Set2* depleted gonads displayed a loss of GSCs, an accumulation of cysts, and a loss of proper egg chamber formation (Figure 1B-D; **Figure 1-Supplement 1A-B’**). The egg chambers that do form contain undifferentiated and differentiating cells marked by spectrosomes and fusomes that fail to develop further (100% in *Set2* RNAi compared to 0% in *nosGAL4*; *p<2.2E-16*, n=50), resulting in females that are infertile. Additionally, *Set2* depleted germ cells had significantly reduced H3K36me3 levels compared to the control, consistent with previous reports (Mukai et al., 2015) (**Figure 1-Supplement 1C-E**).

The accumulation of cyst-like structures upon germline depletion of *Set2* could be due to GSCs that divide but fail to undergo cytokinesis, resulting in GSC cysts, or differentiating cysts that cannot progress further in development (Carreira-Rosario et al., 2016; Mathieu and Huynh, 2017; Sanchez et al., 2016). To discern between these two types of cysts, we stained for pMad, a marker of GSCs. In addition, we crossed a *bam* transcriptional reporter, *bam-GFP*, into Set2 RNAi background and independently assayed for Bam protein (Chen and McKearin, 2003b; Eikenes et al., 2015; Matias et al., 2015). We found that *Set2* RNAi germaria accumulated differentiating cysts, which transcribed and then translated Bam and were pMad negative (Figure 1E-F’; **Figure 1-Supplement 1F-I’**). Thus, Set2 is required in the germline downstream of *bam* to promote the differentiation of Bam expressing cysts into egg chambers.

Although germline depletion of *Set2* leads to both loss of GSCs and accumulation of cysts, here we focus on the cyst accumulation phenotype. Loss of *Set2* results in cysts that do not properly express the oocyte specific protein Orb (Mukai et al., 2015), but loss of Orb does not phenocopy loss of *Set2*, suggesting that Orb downregulation is a consequence of the differentiation defect (Barr et al., 2019; Christerson and McKearin, 1994; Huynh and St Johnston, 2000). Similar to loss of *Set2*, loss of Rbfox1 results in the accumulation of Bam expressing cysts that do not differentiate into proper egg chambers (Carreira-Rosario et al., 2016; Tastan et al., 2010). To test if *Set2* regulates Rbfox1 and Bru expression, we stained separately for Rbfox1 and Bru along with Vasa and 1B1 in control and germline *Set2* depleted ovaries. While control germaria express Rbfox1 robustly in 8-cell cysts, *Set2* depleted germ cells exhibited a significantly lower level of Rbfox1, while somatic levels were unchanged (Figure 1G-I’). Furthermore, Bru levels were reduced and enrichment to the oocyte was ablated in *Set2* RNAi germaria compared to controls (**Figure 1-Supplement 1J-L).** Thus, Set2 is required after Bam expression to promote proper differentiation via Rbfox1 expression.

As germline depletion of *Set2* results in reduced levels of Rbfox1 and Bru, we hypothesized that *Set2* depleted cysts do not properly enter meiosis nor specify an oocyte. To determine if Set2 is required for meiotic progression, we stained control and *Set2* depleted germaria with antibodies against a SC member, Crossover suppressor on 3 of Gowen (C(3)G), and Vasa (Anderson et al., 2005; Page and Hawley, 2001). The control had several C(3)G positive germ cells in 16-cell cysts but only the most posterior germ cell in the egg chamber was marked with C(3)G. In *Set2* germline depleted germaria, the majority of cells displayed perturbed C(3)G expression where an irregular number of cells were C(3)G positive and C(3)G improperly coated DNA and appeared fragmented (Figure 1J-K1’). To determine if the oocyte is properly specified, we stained for the oocyte determinant Egalitarian (Egl), as well as Vasa and 1B1 (Carpenter, 1994; Huynh and St Johnston, 2000; Mach and Lehmann, 1997). While control 16-cell cysts had a single Egl positive cell, *Set2* germline depleted germaria showed diffuse staining of Egl without enrichment in a single cell (**Figure 1-Supplement 1M-N’**). Thus, Set2 is required for proper meiotic progression and oocyte specification.

### MSL3 acts downstream of Set2 to promote entry into meiosis independent of the MSL complex

To identify readers of H3K36me3 that activate transcription downstream of Set2, we screened known Chromodomain (CD) containing proteins, which recognize lysine methylation marks, for loss of function phenotypes that phenocopied Set2 (Allis and Jenuwein, 2016; Bannister et al., 2001; McCarthy et al., 2018a; Nakayama et al., 2001; Navarro-Costa et al.; Yap and Zhou, 2011). Unexpectedly, we identified the H3K36me3 reader MSL3. MSL3 is required in the MSL complex in male flies (Lucchesi and Kuroda, 2015; Samata and Akhtar, 2018), but its role in the female *Drosophila* germline was unknown.

To investigate MSL3 expression in ovaries, we analyzed *msl3* transcript levels at different stages of oogenesis, using RNA-seq libraries that we enriched for GSCs, CBs, cysts, and whole adult ovaries as previously described (McKearin and Ohlstein, 1995; Xie and Spradling, 1998; Zhang et al., 2014). We found that *msl3* mRNA is expressed during oogenesis (**Figure 2-Supplement 1A**). We also examined a fly line expressing GFP tagged MSL3 under endogenous control (Strukov et al., 2011). We stained ovaries from MSL3-GFP flies for GFP and 1B1, and found that MSL3-GFP in the germline was expressed in single cells marked by spectrosomes and early cysts marked by fusomes (Figure 2A, **A’; Figure 2-Supplement 1B**). Thus, MSL3 is expressed in germ cells prior to and during meiotic commitment.

**Figure 2.**
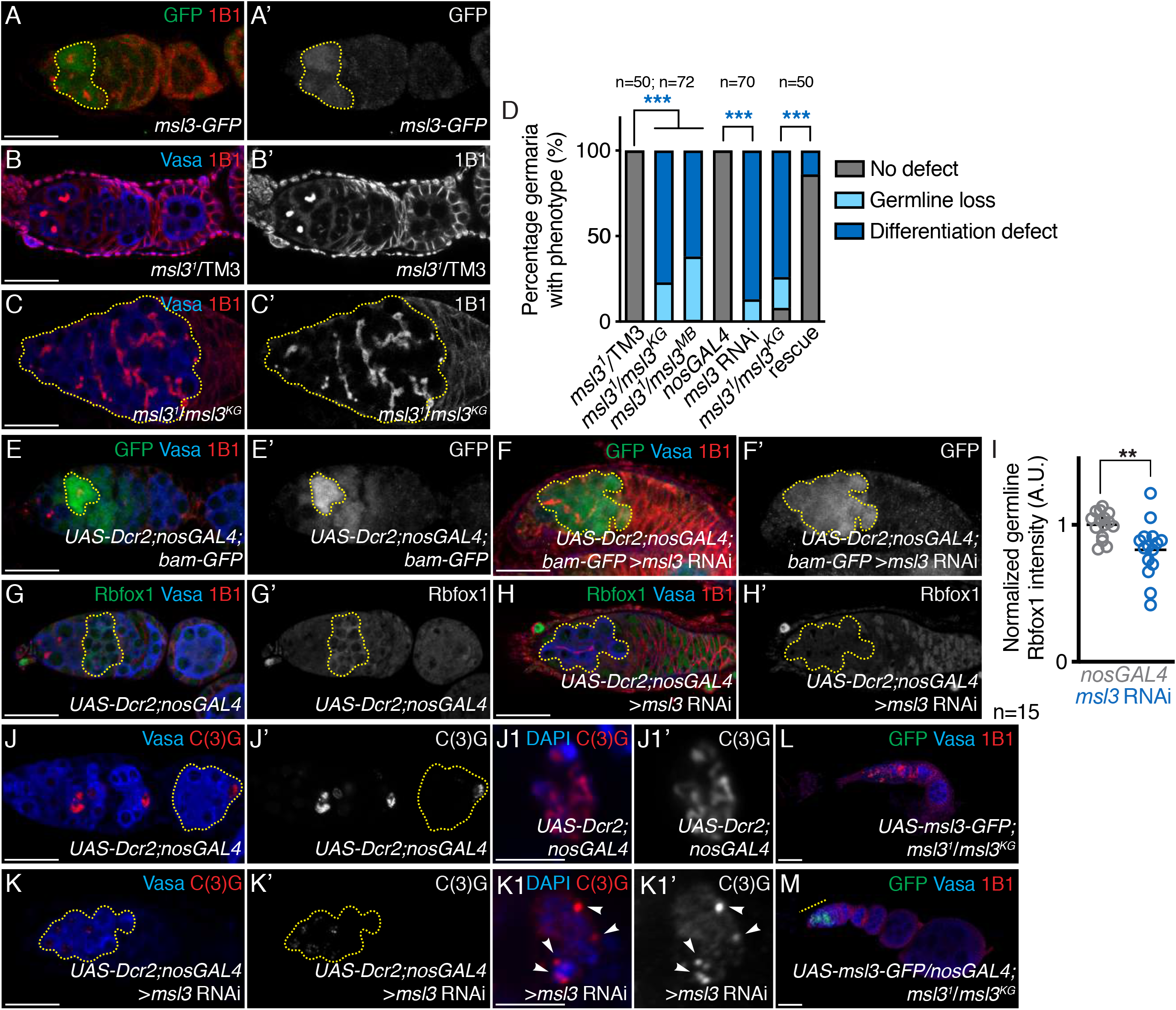
MSL3 is required in the germline for meiotic progression. (A-A’) *msl3-GFP* germarium stained for GFP (green) and 1B1 (red). GFP expression is enriched in single cells and early cysts, showing that MSL3 is expressed in the mitotic and early meiotic stages of oogenesis. GFP channel is shown in A’. (B-B’) Heterozygous control and (C-C’) trans-allelic *msl3* mutant germaria stained for Vasa (blue) and 1B1 (red) shows that *msl3* mutants have irregular cysts (yellow dashed outline) (77% in *msl31/msl3KG* compared to 0% in *msl31* heterozygotes; *p<2.2E-16*, n=50) and germline loss (23% in *msl31/msl3KG* compared to 0% in *msl31* heterozygotes; *p=0.0002*, n=50). 1B1 channel is shown in B’ and C’. Quantitation in (D), statistical analysis performed with Fisher’s exact test on differentiation defect; *** indicates *p<0.001*. (E-E’) Control and (F-F’) germline depleted *msl3* germaria both carrying a *bam-GFP* transgene stained for GFP (green), Vasa (blue), and 1B1 (red) shows that *msl3* germline depletion results in irregular GFP-positive cysts compared to control (yellow dashed outline) (96% in *msl3* RNAi compared to 0% in *nosGAL4*; *p<2.2E-16*, n=50). Statistical analysis performed with Fisher’s exact test. GFP channel is shown in E’ and F’. (G-G’) Control and (H-H’) germline depleted *msl3* germaria stained for Rbfox1 (green), Vasa (blue), and 1B1 (red) shows that *msl3* germline depletion results in decreased levels of Rbfox1 in the germline compared to control (yellow dashed outline) (0.8±0.1 in *msl3* RNAi compared to 1.0±0.1 in *nosGAL4*; *p=0.0037*, n=15). Rbfox1 channel is shown in G’ and H’. Quantitation in (I), statistical analysis performed with Student t-test; ** indicates *p<0.01*. (J-J’) Control and (K-K’) germline depleted *msl3* germaria stained for Vasa (blue) and C(3)G (red) shows that *msl3* germline depletion results in aberrant C(3)G staining compared to control (yellow dashed outline) (100% in *msl3* RNAi compared to 0% in *nosGAL4*; *p<2.2E-16*, n=50) and improper assembly of the synaptonemal complex (white arrows). Statistical analysis performed with Fisher’s exact test. C(3)G channel is shown in J’, J1’, K1, and K1’. (L) Control and (M) germline overexpression of *msl3* in *msl3* mutant germaria stained for GFP (green), Vasa (blue), and 1B1 (red) shows that *msl3* germline overexpression in *msl3* mutants results in reduced frequency of irregular cysts (yellow dashed line) (14% in *msl3* rescue compared to 74% in *msl3* mutant; *p<2.2E-16*, n=50) and germline loss (0% in *msl3* rescue compared to 18% in *msl3* mutant; *p=0.0002*, n=50). Quantitation in (D). Scale bar for J1-J1’ and K1-K1’ is 2 μm, scale bar for all other images is 20 μm.

To verify that MSL3 is required during oogenesis, we examined validated *msl3* mutants (Bachiller and Sánchez, 1989; Sural et al., 2008; Uchida et al., 1981). Indeed, loss of *msl3* lead to cyst accumulation, germline loss, and failure to make egg proper chambers (Figure 2B-D; **Figure 2-Supplement 1C-D’**). Depletion of *msl3* in the germline alone resulted in the accumulation of cysts, phenocopying *msl3* mutants (Figure 2D; **Figure 2-Supplement 2A-B’**). The cysts that accumulate upon *msl3* germline depletion expressed *bam* but were pMad negative and failed to properly express Rbfox1 or Bru, phenocopying *Set2* germline depletion (Figure 2E-I; **Figure 2-Supplement 2C-I).** In addition, the accumulated cysts failed to specify an oocyte, as monitored by Egl, and do not properly express the synaptonemal protein C(3)G (Figure 2J-K1’; **Figure 2-Supplement 2J-K’**). Expression of *msl3* in the germline of *msl3* mutant females was sufficient to rescue the differentiation defect (Figure 2D; L-M). Thus, the H3K36me3 writer, Set2, and the H3K36me3 reader, MSL3, are required in the germline to commit to meiosis.

To determine if MSL3 and Set2 act together to promote oogenesis, we generated flies heterozygous for *Set2* and *msl3*. The germaria of these trans-heterozygous flies displayed severe germline loss compared to single heterozygous controls (**Figure 2-Supplement 1E-G**). Although loss of MSL3 did not affect H3K36me3 levels, loss of *Set2* abolished MSL3 expression (**Figure 2-Supplement 1H-L**). Together, these data suggest that Set2 and MSL3 impinge upon the same developmental pathway(s), with MSL3 acting downstream of Set2 to promote proper meiosis.

In the MSL complex, MSL3 binds to H3K36me3 and helps to recruit MOF, which acetylates H4K16 to promote transcription of the X chromosome (Keller and Akhtar, 2015; Laverty et al., 2010; Lucchesi and Kuroda, 2015). To test if MSL3 functions through the MSL complex in the ovaries, we examined transcript levels of the other MSL complex members (*msl1*, *msl2*, *mof*, *mle*, *roX1*, and *rox2*) (Lucchesi and Kuroda, 2015). We found that *msl1*, *mle*, and *mof* are expressed in ovaries, but *msl2*, *roX1,* and *roX2* are lowly expressed (<1 TPM), consistent with previous reports (Bashaw and Baker, 1997; Meller et al., 1997; Parisi et al., 2004). Additionally, we examined validated *msl1*, *msl2*, and *mle* mutants (Bachiller and Sánchez, 1989b; Belote, 1983; Uchida et al., 1981) and did not observe early oogenesis defects (**Figure 2-Supplement 2L**). Moreover, loss of germline MOF did not result in accumulation of cysts (Sun et al., 2015). Thus, MSL3 functions independently of the MSL complex downstream of Set2 to promote differentiation.

### ATAC complex acts with Set2 and MSL3 to promote meiotic entry

As MSL complex members are either not expressed or not required in the female gonad, we asked if MSL3 cooperates with another HAT-containing complex to regulate cyst differentiation. To identify what HAT works downstream of MSL3, we performed an RNAi screen. We found that members of the Ada2a-containing (ATAC) complex phenocopy loss of Set2 and MSL3 in the germline (Spedale et al., 2012; Suganuma et al., 2008) (Figure 3A-C; **Figure 3-Supplement 1A-H)**. The ATAC complex contains thirteen members, some shared with other complexes, including the HAT Gcn5 (Spedale et al., 2012). Depletion of six members, four of which are specific to the ATAC complex, resulted in accumulation of cysts and germline loss (**Figure 3-Supplement 1A-H**). Of those ATAC complex members, we chose to focus on Negative Cofactor 2β (NC2β), as its defect was highly penetrant but maintained sufficient germline for transcriptomic analysis (see below).

**Figure 3.**
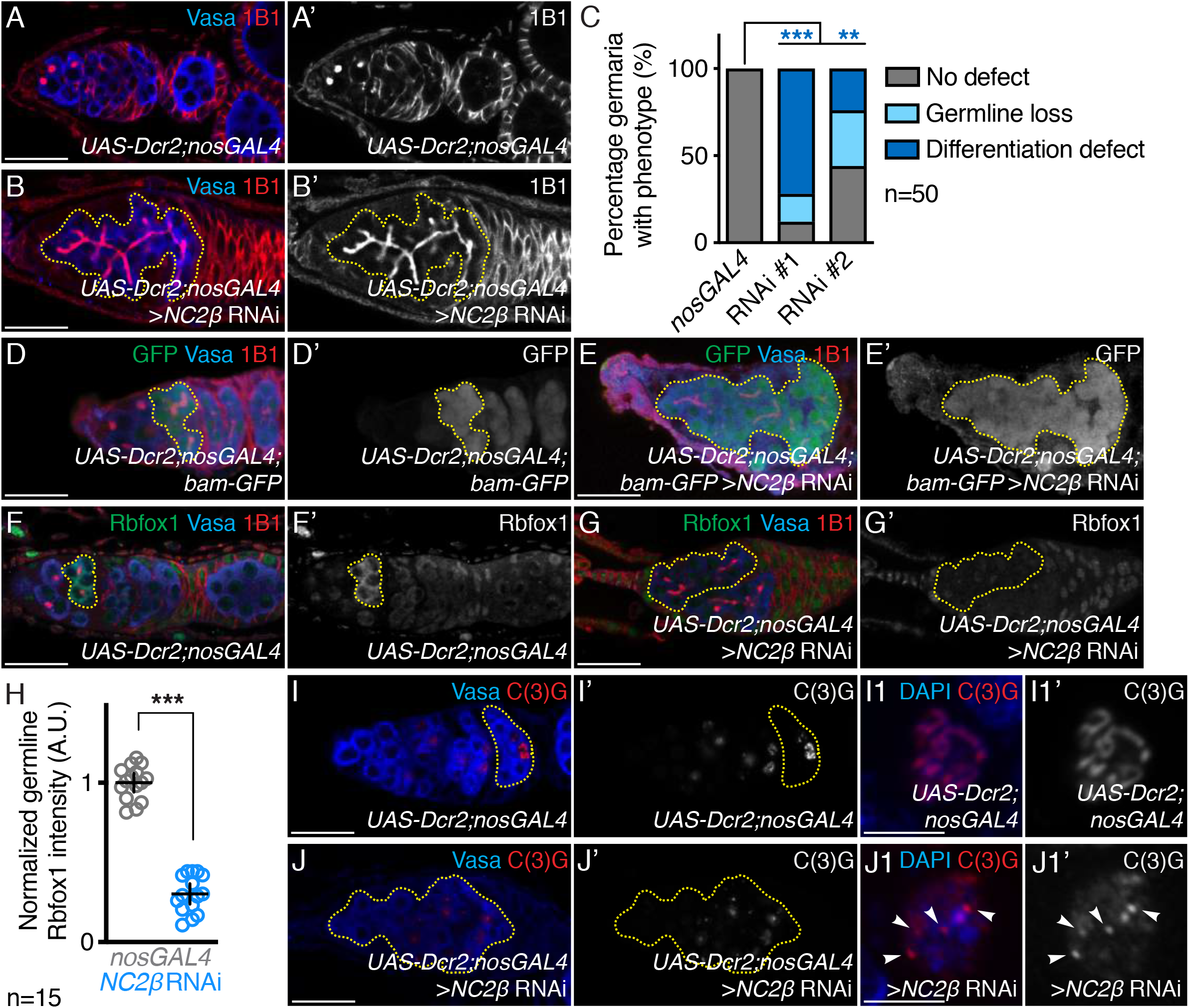
ATAC component, NC2β, is required in the germline for meiotic progression. (A-A’) Control and (B-B’) germline depleted *NC2β* (RNAi line #1) germaria stained for Vasa (blue) and 1B1 (red) shows that *NC2β* germline depletion results in irregular cysts (yellow dashed outline) (72% in *NC2β* RNAi line #1 and 24% in *NC2β* RNAi line #2 compared to 0% *nosGAL4*; *p=9.5E-16* and *p=2.4E-4*, n=50) and germline loss (16% in *NC2β* RNAi line #1 and 21% in *NC2β* RNAi line #2 compared to 0% in *nosGAL4*; *p=0.03* and *p=2.3E-4*, respectively, n=50). 1B1 channel is shown in A’ and B’. Quantitation in (C), statistical analysis performed with Fisher’s exact test on differentiation defect; ** indicates *p<0.01* and *** indicates *p<0.001*. (D-D’) Control and (E-E’) germline depleted *NC2β* germaria both carrying a *bam-GFP* transgene stained for GFP (green), Vasa (blue), and 1B1 (red) shows that *NC2β* germline depletion results in irregular GFP positive cysts compared to control (yellow dashed outline) (64% in *NC2β* RNAi compared to 0% in *nosGAL4*; *p=2.5E-13*, n=50). Statistical analysis performed with Fisher’s exact test. GFP channel is shown in D’ and E’. (F-F’) Control and (G-G’) germline depleted *NC2β* germaria stained for Rbfox1 (green), Vasa (blue), and 1B1 (red) shows that *NC2β* germline depletion results in decreased levels of Rbfox1 in the germline compared to control (yellow dashed outline) (0.3±0.1 in *NC2β* RNAi compared to 1.0±0.1 in *nosGAL4*; *p<0.0001*, n=50). Rbfox1 channel is shown in F’ and G’. Quantitation in (H), statistical analysis performed with Student t-test; *** indicates *p<0.001*. (I-I’ and I1-I1’) Control and (J-J’ and J1-J1’) germline depleted *NC2β* germaria stained for Vasa (blue) and C(3)G (red) shows that *NC2β* germline depletion results in aberrant C(3)G staining compared to control (yellow dashed outline and white arrows) (75% in *NC2β* RNAi compared to 0% in *nosGAL4*; *p<2.2E-16*, n=50) and improper assembly of the synaptonemal complex (white arrows). Statistical analysis performed with Fisher’s exact test. C(3)G channel is shown in I’, I1’, J’, and J1’. Scale bar for I1-I1’ and J1-J1’ is 2 μm, scale bar for all other images is 20 μm.

Loss of NC2β in the germline led to GSC loss and accumulation of cysts-like structures that were marked by fusomes (Figure 3A-C; **Figure 3-Supplement 1A-B’**). These cysts expressed Bam, did not contain pMad positive cells or properly express Rbfox1 or Bru (Figure 3D-H; **Figure 3-Supplement 2A-G**). In addition, loss of *NC2β* leads to loss of meiotic progression and oocyte specification as monitored by C(3)G localization and Egl respectively (Figure 3I-J1’; **Figure 3-Supplement 2H-I’**). These data suggest that the ATAC complex, like Set2 and MSL3, is required for commitment to a meiotic program as well as oocyte specification.

As components of ATAC complex phenocopy loss of *Set2* and *msl3*, we asked whether the ATAC complex may acts together with MSL3 to promote meiotic entry. To test this, we stained for H3K36me3 in *NC2β* RNAi flies and found that H3K36me3 levels were unaltered (**Figure 3-Supplement 2J-L**). In addition, we made use of a mutant of the active HAT in the ATAC complex, Atac2, as there were no available *NC2β* mutants available. We generated flies heterozygous for both *Atac2* and *msl3*, and found that their germaria had severe oogenesis defects compared to the single heterozygous controls (**Figure 3-Supplement 2M-O**). Thus, the ATAC complex works downstream of Set2, and *Atac2* genetically interacts with *msl3*. Taken together, our data suggest that Set2, MSL3, and ATAC complex impinge upon the same developmental pathway(s) to regulate meiotic progression in the *Drosophila* female germline.

### Set2, MSL3, and NC2β promote transcription of the ribosomal protein paralog *RpS19b*

To determine how Set2, MSL3, and ATAC promote meiotic commitment, we compared the transcriptomes of *Set2*, *msl3*, and *NC2β* germline depleted ovaries with to a developmental control that accumulates cysts. To enrich for cysts we induced *bam* expression under control of a *heat-shock* (*hs*) promoter in the background of germaria depleted for *bam* (*bam* RNAi;*hs-bam*) (Ohlstein and McKearin, 1997; Zhang et al., 2014). We found 662 significantly downregulated RNAs, whereas 65 RNAs were upregulated in *Set2* depleted germaria compared to *bam* RNAi;*hs-bam* ovaries (Fold Change (FC)=4; False discovery rate (FDR)=0.05) (Figure 4A). There were 283 significantly downregulated RNAs and 302 significantly upregulated RNAs in *msl3* RNAi compared to *bam* RNAi;*hs-bam* (Figure 4A’). Lastly, there were 466 RNAs significantly downregulated and 277 upregulated, in *NC2β* RNAi compared to the developmental control (Figure 4A’’). Of those transcripts that were differentially expressed in *Set2*, *msl3*, and *NC2β* depleted germ cells compared to *bam* RNAi;*hs-bam* control there were 29 shared RNAs that were downregulated (Figure 4B) and 11 shared RNAs that were upregulated. As these transcriptional regulators are known to promote transcription, we focused on the downregulated RNAs.

**Figure 4.**
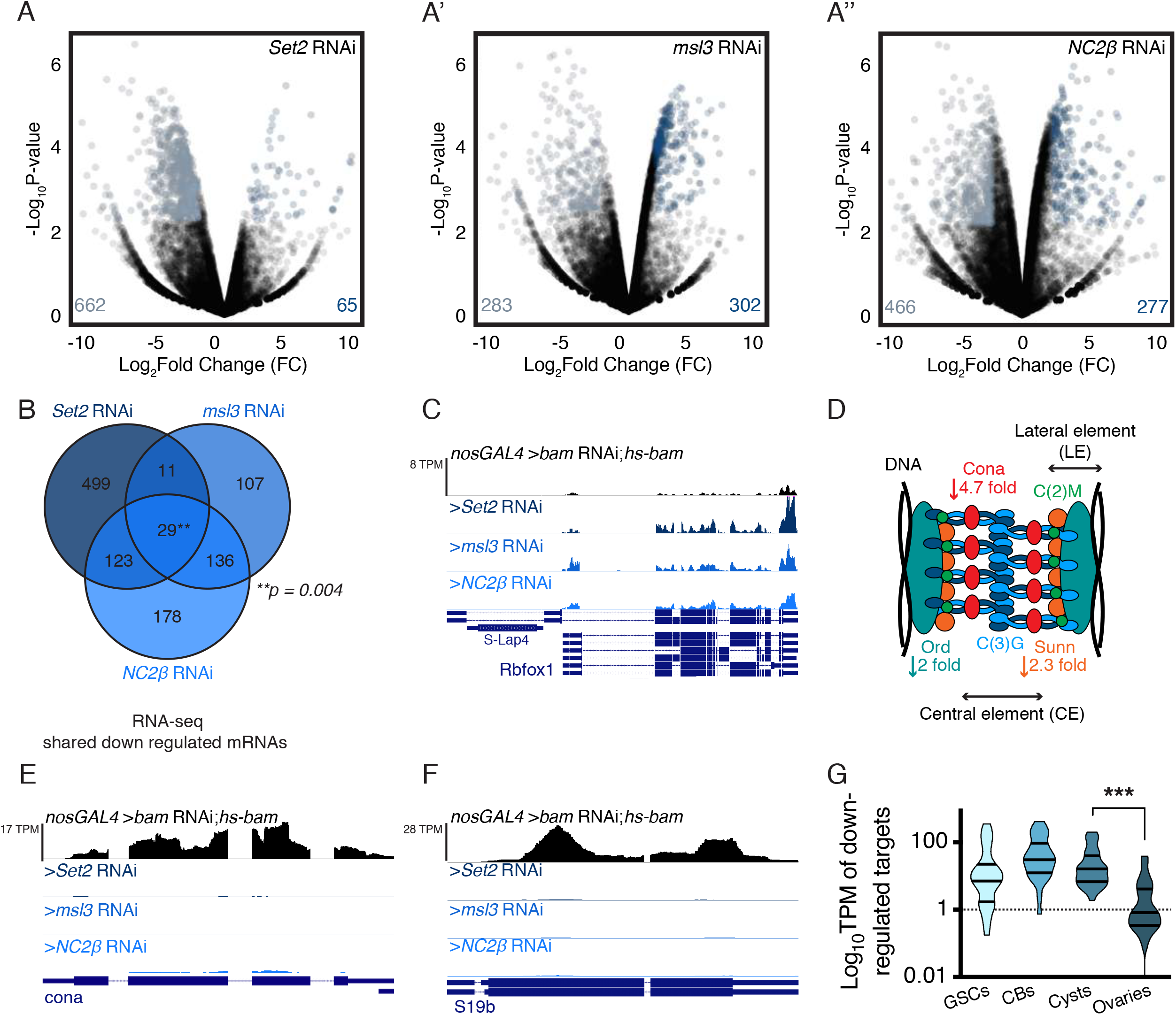
Set2, MSL3, and ATAC complex regulate mRNA levels of recombination machinery components, but not *Rbfox1*. (A-A’’) Volcano plots of –Log10P-value vs. Log2Fold Change (FC) of (A) *Set2*, (A’) *msl3*, and (A’’) *NC2β* germline depleted ovaries compared to *bam* RNAi;*hs-bam*. Light blue dots represent significantly downregulated transcripts and dark blue dots represent significantly upregulated transcripts in *Set2*, *msl3*, and *NC2β* RNAi ovaries compared with *bam* RNAi;*hs-bam* ovaries (FDR = 0.05). Genes with four-fold or higher change were considered significant. (B) Venn diagram of downregulated genes from RNA-seq of *Set2*, *msl3*, and *NC2β* germline depleted ovaries compared to *bam* RNAi;*hs-bam*. 29 targets are shared between *Set2*, *msl3*, and *NC2β* RNAi, suggesting that while Set2, MSL3, and ATAC function independently they also co-regulate a small population of genes. (C) RNA-seq track showing that *Rbfox1* is not reduced upon germline depletion of *Set2*, *msl3*, and *NC2β*. All tracks are set to scale to 8 TPM. (D) A structural model of the SC where the lateral element (LE), consisting of proteins such as Ord (teal), Sunn (orange), and C(2)M (green) assemble along DNA. The central region (CR) consists of transverse elements such as Cona (red) and C(3)G (light and dark blue) to stabilize the complex and promote recombination. Down arrows denote fold downregulation of SC components in depleted ovaries. (E) RNA-seq track showing that *cona* is reduced upon germline depletion of *Set2*, *msl3*, and *NC2β*. All tracks are set to scale to 17 TPM. (F) RNA-seq track showing that *RpS19b* is reduced upon germline depletion of *Set2*, *msl3*, and *NC2β*. All tracks are set to scale to 28 TPM. (G) Violin plot of mRNA levels of the 29 shared downregulated targets in ovaries enriched for GSCs, CBs, cysts, and whole ovaries, showing that the shared targets are most highly enriched in CBs and cyst stages, that then tapers off in whole ovaries (41.2±15.1 in single cells, 76.2±19.1 in, 35.6±9.5 in cyst, and 4.2±1.6 in whole *nosGAL4* ovaries; *p=0.009* for whole ovaries compared to cysts). Statistical analysis performed with one-way ANOVA; *** indicates *p<0.001*.

Interestingly, although Rbfox1 protein is not properly expressed upon loss of Set2, MSL3, and NC2β, *Rbfox1* mRNA was not among the shared downregulated RNAs (Figure 4C). We verified that *Rbfox1* mRNA was present in germline of *msl3* depleted ovaries by *in situ* hybridization (**Figure 4-Supplement 1A-B’**). Thus, Set2, MSL3, and ATAC do not regulate transcription of *Rbfox1* mRNA to promote meiotic commitment. In contrast, we found that several SC member genes were among the shared downregulated genes, including *orientation disruptor* (*ord*), *sisters unbound* (*sunn*), and *corona* (*cona*) (Hughes et al., 2018) (Figure 4D-E; **Figure 4-Supplement 1C-D**). To validate the loss of SC components, we crossed an Ord-GFP line (Balicky et al., 2002) into *msl3* mutants and found that *msl3* mutant ovaries had both lower GFP levels as well as mislocalized Ord compared to controls (**Figure 4-Supplement 1E-F’**). The shared downregulated targets also included 11 candidate genes (CGs) of unknown function, and the ribosomal protein paralog, *RpS19b*, but not *RpS19a* (Figure 4B, F; **Figure 4-Supplement 1G-J’**).

We hypothesized that MSL3 and its directly regulated downstream targets would be expressed at the same stages, from GSCs until the cyst stages. To test this hypothesis, we analyzed mRNA levels of the 29 targets in RNA-seq libraries enriched for either GSCs, CBs, cysts, or unenriched wild type ovaries. Indeed, transcript levels overlap with MSL3 expression and then dropped off (Figure 4G). Taken together, these data suggest that the Set2, MSL3, and ATAC axis regulates transcription of SC components and *RpS19b,* but not *Rbfox1*, during GSC differentiation.

### RpS19b is a germline enriched ribosomal protein required for Rbfox1 translation

RpS19b is a ribosomal protein and is one of two RpS19 paralogs, RpS19a and RpS19b in *Drosophila* (Marygold et al., 2007; Shigenobu et al., 2006). These two paralogs are ∼80% similar (Sayers et al., 2012, FlyBase DIOPT v7.1). Humans only have one version of RpS19 (hRpS19/hS19). In humans, reduced expression of RpS19 leads to ribosomopathies due to decreased translation of specific mRNAs, such as the transcription factor GATA1 in the case of Diamond-Blackfan anemia (DBA) (Draptchinskaia et al., 1999; Gazda et al., 2004; Khajuria et al., 2018; Ludwig et al., 2014; Willig et al., 2000).

Given that loss of Set2, MSL3, and NC2β decreased Rbfox1 protein levels without affecting *Rbfox1* mRNA levels, we hypothesized that reduced RpS19b expression resulted in decreased translation of *Rbfox1* mRNA. If RpS19b is required for translation of Rbfox1, then RpS19b and Rbfox1 protein expression should overlap. We examined lines expressing RpS19b-GFP and RpS19a-HA from their endogenous promoters. RpS19b-GFP was germline enriched while RpS19a-HA was expressed in both the germline and soma of gonad (Figure 5A-A1; **Figure 5-Supplement 1A-B**). In the germline, RpS19b-GFP was expressed at high levels in single cells and gradually decreased in cyst stages, which overlapped with the protein expression of MSL3 and Rbfox1 (Figure 5B).

**Figure 5.**
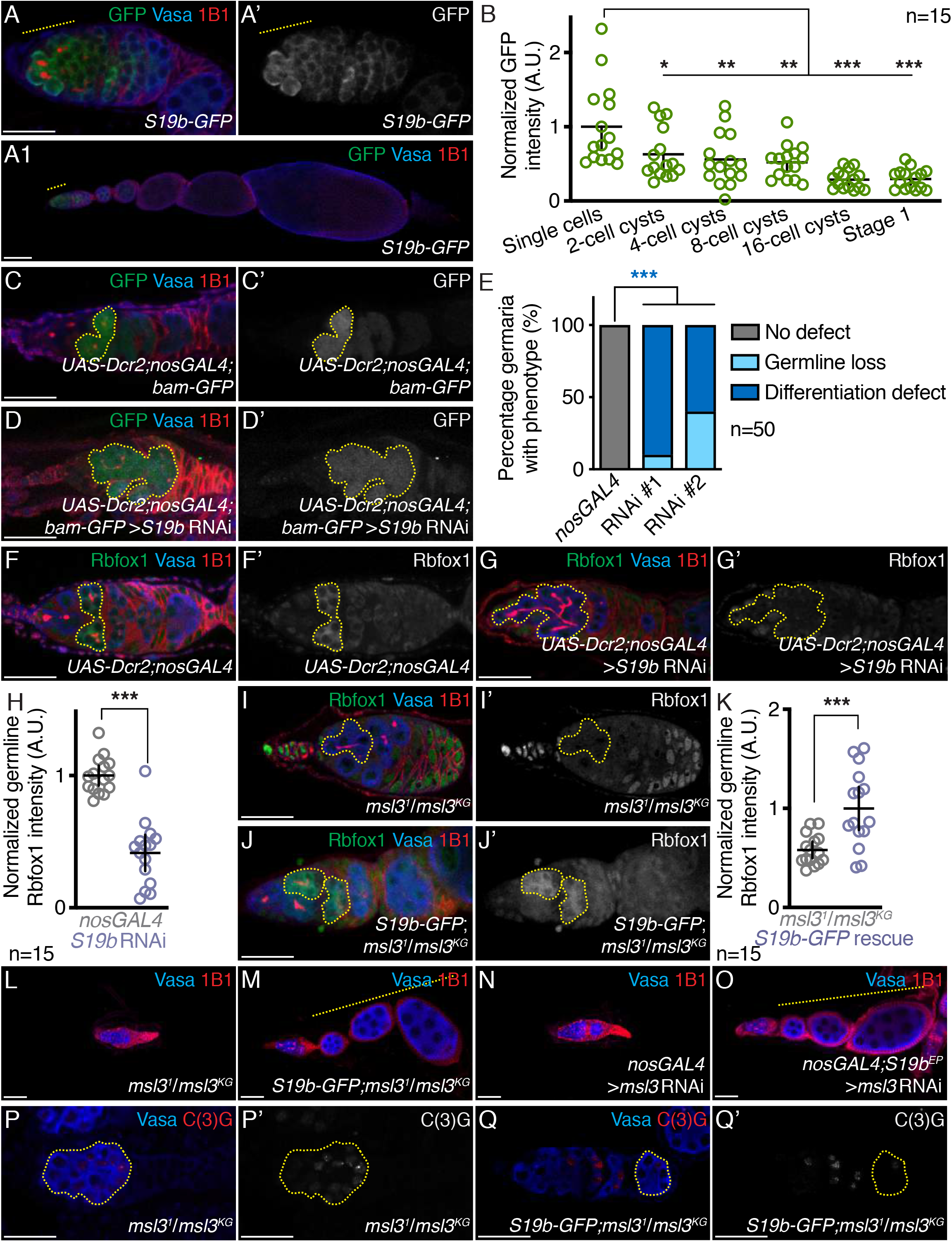
RpS19b, a germline enriched paralog, is expressed in the mitotic and early meiotic stages and is required for Rbfox1 expression. (A-A’) *RpS19b-GFP* germarium and (A1) ovariole stained for GFP (green), Vasa (blue), and 1B1 (red). GFP is expressed higher in single cells in the germarium, decreases in the cyst stages, and then tapers off upon stage 1 formation (1.0±0.1 in single cells, 0.6±0.1 in 2-cell cyst, 0.6±0.1 in 4-cell cyst, 0.5±0.1 in 8-cell cyst, 0.3±0.1 in 16-cell cyst, and 0.3±0.1 in stage 1 egg chamber; *p=0.0284* for 2-cell cyst, *p=0.0047* for 4-cell cyst, *p=0.0017* for 8-cell cyst, *p<0.0001* for 16-cell cyst and stage 1 egg chamber, compared to single cells, n=15). GFP channel is shown in A’. Quantitation in (B), statistical analysis performed with one-way ANOVA; * indicates *p<0.05*, ** indicates *p<0.01*, and *** indicates *p<0.001*. (C-C’) Control and (D-D’) germline depleted *RpS19b* germaria both carrying a *bam-GFP* transgene stained for GFP (green), Vasa (blue), and 1B1 (red) shows that *RpS19b* germline depletion results in irregular GFP positive germ cells compared to control (yellow dashed outline) (64% in *RpS19b* RNAi compared to 0% in *nosGAL4*; *p=2.5E-13*, n=50). Statistical analysis performed with Fisher’s exact test. GFP channel is shown in C’ and D’. Quantitation in (E), statistical analysis performed with Fisher’s exact test on differentiation defect; *** indicates *p<0.001*. (F-F’) Control and (G-G’) germline depleted *RpS19b* germaria stained for Rbfox1 (green), Vasa (blue), and 1B1 (red) shows that *RpS19b* germline depletion results in decreased levels of Rbfox1 in the germline compared to control (yellow dashed outline) (0.4±0.2 in *RpS19b* RNAi compared to 1.0±0.1 in *nosGAL4*; *p<0.0001*, n=15). Rbfox1 channel is shown in F’ and G’. Quantitation in (H), statistical analysis performed with Student t-test; *** indicates *p<0.001*. (I-I’) Control and (J-J’) *RpS19b-GFP* rescue germaria stained for Rbfox1 (green), Vasa (blue), and 1B1 (red) shows that addition of *RpS19b-GFP* to *msl3* mutants results in increased levels of Rbfox1 expression compared to control (1.0±0.4 in rescue compared to 0.6±0.2 in *msl31/msl3KG*; *p<0.0006*, n=15). Rbfox1 channel is shown in I’ and J’. Quantitation in (K), statistical analysis performed with Student t-test; *** indicates *p<0.001*. (L) Control and (M) *RpS19b-GFP* rescue ovarioles stained for Vasa (blue) and 1B1 (red) shows that addition of *RpS19b-GFP* to *msl3* mutants results in an increased frequency of spectrosomes and cysts (92% in *RpS19b-GFP* rescue compared to 4% in *msl31/msl3KG*; *p<2.2E-16*, n=50) and subsequent egg chambers compared to control (yellow dashed outline) (98% in *RpS19b-GFP* rescue compared to 16% in *msl31/msl3KG*; *p<2.2E-16*, n=50). Statistical analysis performed with Fisher’s exact test. (N) Control and (O) *RpS19bEP* rescue ovarioles stained for Vasa (blue) and 1B1 (red) shows that expression of *RpS19bEP* in *msl3* germline depletion ovaries results in an increased frequency of spectrosomes and cysts (90% in *RpS19bEP* rescue compared to 0% in *msl3* RNAi; *p<2.2E-16*, n=50) and subsequent egg chambers compared to control (yellow dashed outline) (100% in *RpS19bEP* rescue compared to 4% in *msl3* RNAi; *p<2.2E-16*, n=50). Statistical analysis performed with Fisher’s exact test. (P-P’) Control and (Q-Q’) *RpS19b-GFP* rescue germaria stained for Vasa (blue) and C(3)G (red) shows that rescue and control germaria have aberrant C(3)G expression (yellow dashed outline) (100% in *RpS19b-GFP* rescue compared to 100% in *msl31/msl3KG*; *p=1*, n=50). Addition of *RpS19b-GFP* does not rescue egg laying defects (38 eggs/female in *RpS19b-GFP*, 32 eggs/female in *msl31* heterozygote, 101 eggs/female in *msl3KG* heterozygote compared to 0 eggs/female in *msl3KG/msl31* and rescue; *p<0.0001* for all, n=4). Statistical analysis performed with Fisher’s exact test. C(3)G channel is shown in P’ and Q’. Scale bar for all images is 20 μm.

If RpS19b acts downstream of MSL3 to promote translation of *Rbfox1* mRNA, then loss of RpS19b should phenocopy *msl3* mutants, with reduced Rbfox1 protein levels. We used RNAi to specifically deplete *RpS19b* but not *RpS19a* in the germline (**Figure 5-Supplement 1C-F’**) and found that *RpS19b* depleted germaria accumulated *bam*-positive cysts that lack Rbfox1 protein (**Figure 5-Supplement 1G-H’;** Figure 5C-H). We next asked whether addition of *RpS19b* could rescue the differentiation defect upon loss of *msl3*. We found that addition of one copy of *RpS19b-GFP* in *msl3* mutant flies rescued the early cyst defect, including Rbfox1 expression, and lead to egg chamber formation (Figure 5I-M). In addition, overexpression of *RpS19b* via an *EP* line could also rescue the differentiation defect upon germline depletion of *msl3,* leading to egg chamber formation (Figure 5N-O). Thus, our data suggest that MSL3 promotes the expression of RpS19b and thus *Rbfox1* translation and proper entry into meiosis.

Our model predicts that the MSL3-mediated regulation of SC members is independent of Rbfox1 protein expression. To test this model, we examined the localization of the SC component C(3)G in *msl3* mutants that express RpS19b (Anderson et al., 2005; Page and Hawley, 2001). We found that while *msl3* mutants with restored RpS19b expression make egg chambers, C(3)G does not properly localize to the oocyte nucleus in egg chambers and the females were infertile (Figure 5P-Q’). Thus, RpS19b is not involved in MSL3-mediated regulation of SC members to promote recombination during meiosis.

### RpS19 levels, not paralog specificity, are critical for meiotic progression

We generated a CRISPR null mutant of *RpS19b* (*RpS19b^CRISPR^*) that are viable and unexpectedly did not display any oogenesis defects (**Figure 5-Supplement 1I-K**), unlike homozygous *RpS19a* mutants, which are lethal (Shigenobu et al., 2006). Studies in organisms including zebrafish have reported transcriptional compensation in mutants, but not in gene depletion using RNA interference methods (El-Brolosy et al., 2019). To determine if there are transcriptional changes in *RpS19b^CRISPR^* mutants, we performed RNA-seq of CB enriched ovaries utilizing *bam* RNAi and compared it to *RpS19b^CRISPR^* in *bam* depleted background. We enriched for undifferentiated stages as *RpS19b* is primarily expressed only up to the cyst stages. We found that loss of *RpS19b* resulted in 672 downregulated genes and 2,030 upregulated genes with 6-fold downregulation of *RpS19b* but no increase in *RpS19a* levels (1592 TPM in *bam* RNAi;*RpS19b^CRISPR^* compared to 1688 TPM in *bam* RNAi) (**Figure 5-Supplement 1L**). Intriguingly, a translation initiation factor, *eukaryotic translation initiation factor 4B* (*eIF4B*), was upregulated more than 8-fold in *RpS19b* mutants (5.7 TPM in *bam* RNAi;*RpS19b^CRISPR^* compared to 0.67 TPM in *bam* RNAi) suggesting modulation of translation machinery. We then asked if *RpS19b* mutants have proper development because they translate increased levels of RpS19a protein. Using an RpS19 antibody that detects both paralogs, we found that levels of RpS19 were not downregulated in mutant compared to control gonads (**Figure 5-Supplement 1M-P**). Furthermore, germline depletion of *RpS19a* in *RpS19b^CRISPR^* mutants results in complete loss of the germline, compared to no defect in homozygous *RpS19b^CRISPR^* mutants or accumulation of cysts in *RpS19a* depletion alone (**Figure 5-Supplement 1Q-R**). In contrast, *RpS19b* depletion in *RpS19b^CRISPR^* mutants did not have a defect (**Figure 5-Supplement S-T**). Thus, loss of RpS19b can be compensated by increased levels of RpS19a, via yet unknown mechanisms.

Our data suggests that RpS19b expression acts to increase the levels of RpS19, which then promotes expression of Rbfox1. To test this, we depleted *RpS19a* from the germline and found that the germaria accumulate *bam*-positive cysts that have significantly reduced levels of Rbfox1 (**Figure 5-Supplement 2A-L’**). In addition, ectopic expression of RpS19a-HA in *msl3* depleted ovaries restored Rbfox1 protein expression and egg chamber formation but females were infertile and had perturbed C(3)G localization (**Figure 5-Supplement 2M-R**). Furthermore, expression of human RpS19 in the germline of *msl3* depleted germaria also rescued the cyst accumulation phenotype giving rise to egg chambers (**Figure 5-Supplement 2S-T1**). Thus, our data taken together suggests that proper dosage of RpS19 is essential for translation of Rbfox1 protein, that then promotes transition to a meiotic cell fate, and egg chamber formation in *Drosophila*.

### RpS19 promotes Rbfox1 translation in the germline

As proper RpS19 levels are required for Rbfox1 protein expression, we hypothesized that RpS19 regulates translation of *Rbfox1*. To test this, we performed polysome profiling followed by western blot analysis using ovaries enriched for undifferentiated germ cells (*bam* RNAi), as well as whole ovaries. While RpS19a-HA is present in polysome fractions in whole ovaries, RpS19b-GFP appeared to be preferentially enriched in actively translating ribosomes early in oogenesis, consistent with its expression pattern (Figure 6A-B’). To test if RpS19 paralogs affect translation in cysts, we pulsed gonads with a puromycin analog, O-propargyl-puromycin (OPP), that is incorporated into translated peptides and can be detected using Click-chemistry (Sanchez et al., 2016). We found that cysts that accumulate upon the loss of RpS19a and RpS19b have decreased translation compared to cysts of control ovaries (Figure 6C-F).

**Figure 6.**
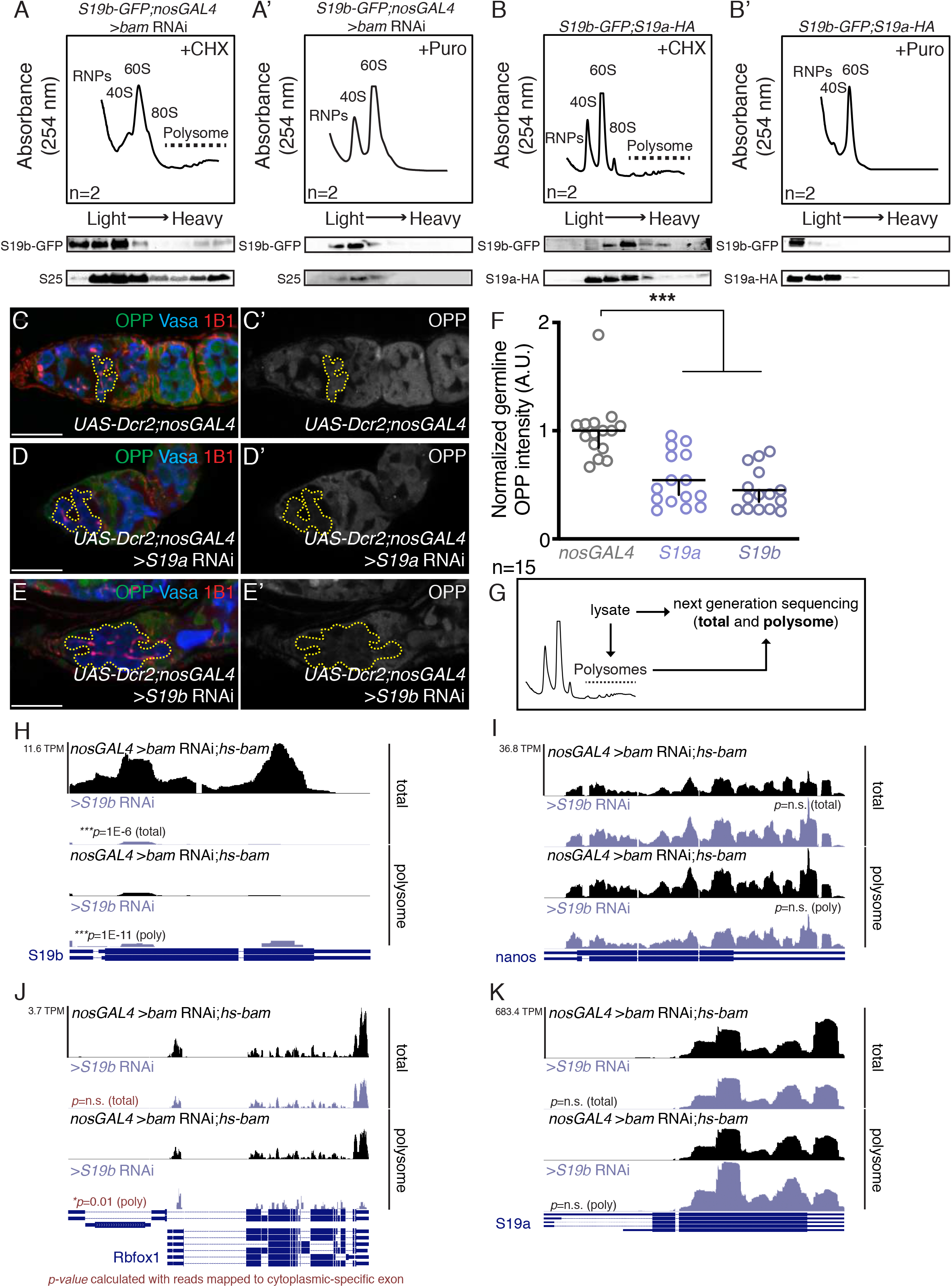
RpS19 paralogs are incorporated into the ribosome and RpS19 levels affect translation, including translation of Rbfox1. (A) Top: Polysome profiles of *RpS19b-GFP;nosGAL4* >*bam* RNAi ovaries treated with cycloheximide (CHX) or (A’) puromycin and fractionated. Polysome profiles show that peaks are present in the polysome (heavy) fractions and are ablated upon puromycin mediated dissociation. Bottom: Western blot analysis of polysome fractionated *RpS19b-GFP* CB enriched ovaries treated with (A) cycloheximide (CHX) or (A’) puromycin and fractionated. Blots were stained for GFP (top) and RpS25 (bottom), showing RpS19b and RpS25 bands in heavy fractions in CHX-treated samples that are absent in puromycin treated samples. (B) Top: Polysome profiles of *RpS19b-GFP;RpS19a-HA* whole ovaries treated with cycloheximide (CHX) or (B’) puromycin and fractionated. Polysome profiles show that peaks are present in the polysome (heavy) fractions and are ablated upon puromycin mediated dissociation. Bottom: Western blot analysis of polysome fractionated *RpS19b-GFP;RpS19a-HA* whole ovaries treated with (B) cycloheximide (CHX) or (B’) puromycin and fractionated. Blots were stained for HA (top) and GFP (bottom), showing RpS19a and RpS19b bands in heavy fractions in CHX-treated samples that are absent in puromycin treated samples. (C-C’) Control, (D-D’) germline depleted *RpS19a* and (E-E’) *RpS19b* germaria pulsed with OPP (green) and stained for Vasa (blue) and 1B1 (red) shows that *RpS19a* and *RpS19b* germline depletion results in decreased OPP compared to control (0.5±0.2 in *RpS19a* RNAi and 0.4±0.2 in *RpS19b* RNAi compared to 1.0±0.3 in *nosGAL4*; *p<0.0001* for both, n=15). OPP channel is shown in C’, D’, and E’. Quantitation in (F), statistical analysis performed with one-way ANOVA; *** indicates *p<0.001*. (G) A schematic of the experimental approach to polysome-seq where RNA is extracted (total) with polysome fractionation (polysome) followed by next generation sequencing. (H) RNA-seq track of total (top) and polysome (bottom) showing that *RpS19b* is reduced upon germline depletion of *RpS19b* (purple) compared to control (black) (total: Log2FC=-4.1, *p=1E-6*, n=2 and polysome: Log2FC=-4.5; *p=1E-11*, n=2). All tracks are set to scale to 11.6 TPM. Statistical analysis performed with Student t-test; *** indicates *p<0.001*. (I) RNA-seq track of total (top) and polysome (bottom) showing that *nanos* and amount of germline is not reduced upon germline depletion of *RpS19b* (purple) compared to control (black) (total: Log2FC=0.4; *p=0.4*, n=2 and polysome: Log2FC=0.3; *p=0.7*, n=2). All tracks are set to scale to 36.8 TPM. Statistical analysis performed with Student t-test; “n.s.” indicates *p>0.5*. (J) RNA-seq track of total (top) and polysome (bottom) showing that cytoplasmic *Rbfox1* is reduced in polysome fractions upon germline depletion of *RpS19b* (purple) compared to control (black) (total: *p=0.2*, n=2 and polysome: *p=0.01*, n=2). All tracks are set to scale to 3.7 TPM. Statistical analysis performed with Student t-test; “n.s.” indicates *p>0.5* and * indicates *p<0.05*. (K) RNA-seq track of total (top) and polysome (bottom) showing that *RpS19a* is not reduced upon germline depletion of *RpS19b* (purple) compared to control (black) (total: Log2FC=-0.4; *p=0.4*, n=2 and polysome: Log2FC=-0.1; *p=0.9*, n=2). All tracks are set to scale to 683.4 TPM. Statistical analysis performed with Student t-test; “n.s.” indicates *p>0.5*. Scale bar for all images is 20 μm.

To directly test whether RpS19b is required for *Rbfox1* translation we then performed polysome-seq on germaria depleted of *RpS19b* compared to control germaria enriched for cysts using *bam RNAi*;*hs-bam* (Figure 6G-H). Depletion of germline *RpS19b* did not significantly affect the translation efficiency of germline specific mRNA, *nanos,* but there was a reduction of *Rbfox1* mRNA translation efficiency, compared to control (Figure 6I-J). Additionally, depletion of *RpS19b* using RNAi, did not reduce the levels or translation efficiency of *RpS19a* (Figure 6K). Taken together, our data suggest that there is an increased expression of RpS19 during early development that is required for translation of *Rbfox1* mRNA.

### MSL3 is expressed during meiotic stages of mouse spermatogenesis

As Set2, MSL3, and ATAC complex are conserved in mammals, we hypothesized that this transcriptional axis could also regulate meiotic entry in mammals. Set2 and ATAC are general transcriptional regulators and present in most tissues, therefore we asked if MSL3 is differentially expressed in the male gonad, which is easily accessible and because of germline stem cells, has ongoing meiosis (Guelman et al., 2009; Li et al., 2016). In male mice, undifferentiated Type A spermatogonial stem cells (SSCs) reside proximal to the basement membrane of the seminiferous tubules where they self-renew and divide (Boyle et al., 2007; Hess and de Franca, 2008; Oatley and Brinster, 2008; Ohta et al., 2003; Ryu et al., 2006). These spermatogonia are maintained by support cells called Sertoli cells (Hess and R. França, 2005). Upon retinoic acid (RA) signaling, Type B spermatogonia express markers such as KIT receptor (cKIT) and STRA8, differentiate and undergo meiosis to give rise to spermatocytes (SPCs), spermatids, and spermatozoa (Busada et al., 2015; Endo et al., 2015; Schrans-Stassen et al., 1999; Zhou et al., 2008). To examine spermatogenesis, we stained post-pubertal gonads for the differentiation markers cKIT and STRA8. We observed STRA8 positive germ cells co-localized with cKIT positive SSCs and primary spermatocytes that have reached the pre-leptotene phase of meiosis I, as previously reported (Busada et al., 2015; van Pelt et al., 1995) (Figure 7A-B). To examine the spatiotemporal regulation of MSL3 during spermatogenesis, we stained for cKIT and MSL3. We found that MSL3 forms nuclear speckles in cKIT positive spermatogonia and is nuclear in spermatocytes that are undergoing meiosis (Figure 7C-E). To interrogate where in the nucleus this MSL3 nuclear foci forms in SSCs we co-stained with Synaptonemal complex protein 3 (SYCP3) and DAPI. We found that while chromosomes were coated with SYCP3, MSL3 foci were restricted to the non-recombining chromosomes (Figure 7F-F’’). Taken together, we find that MSL3 is nuclear in cells undergoing meiosis during mouse spermatogenesis. Thus, MSL3 is expressed downstream of STRA8 in meiotic cells during spermatogenesis.

**Figure 7.**
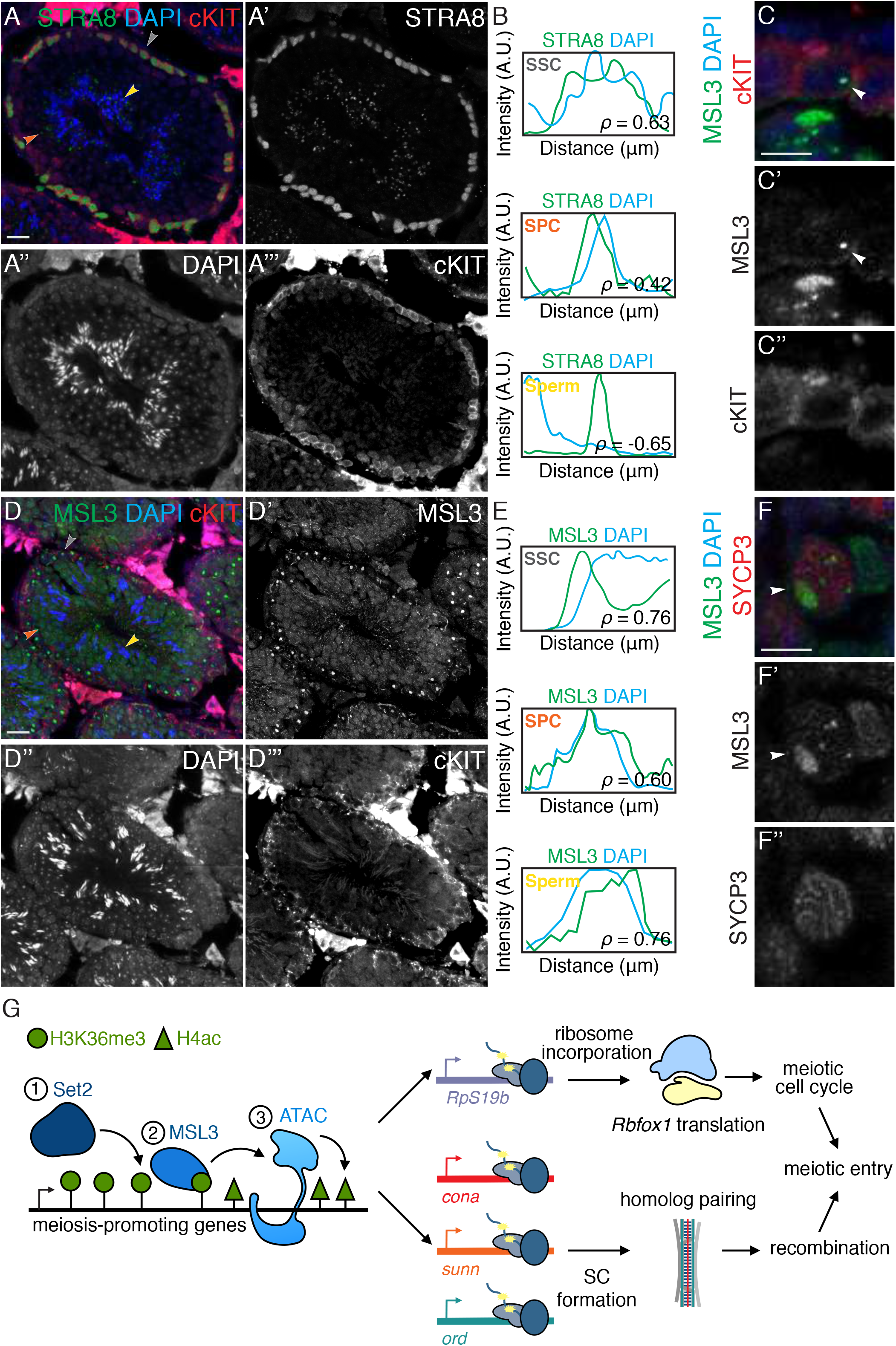
MSL3 is expressed during meiosis in the mouse male gonad. (A-A’’’) Adult gonads stained for STRA8 (green), DAPI (blue), and cKIT (red). cKIT-positive SSCs and SPCs co-stain for STRA8 (gray and orange arrows), shows that STRA8 marks differentiating and pre-leptotene germ cells. Statistical analysis performed with Spearman’s correlation test. STRA8 channel is shown in A’, DAPI channel is shown in A’’, and cKIT channel is shown in A’’’. (B) Correlation plots of STRA8 (green) and DAPI (blue) showing positive correlation between STRA8 and DNA in SSCs (top, gray arrow, ρ=0.63) and SPCs (middle, orange arrow, ρ=0.47), but a negative correlation with spermatozoa (bottom, yellow arrow, ρ=-0.65). (C-C’’) SSC stained for MSL3 (green), DAPI (blue), and cKIT (red) shows that differentiating SSC have MSL3 foci. MSL3 channel is shown in C’ and cKIT channel is shown in C’’. (D-D’’) Adult gonads stained for MSL3 (green), DAPI (blue), and cKIT (red) shows that MSL3 forms nuclear foci in cKIT positive SSCs and then accumulates as smaller foci, coating nuclei as spermatogenesis proceeds. Statistical analysis performed with Spearman’s correlation test. MSL3 channel is shown in D’, DAPI channel is shown in D’’, and cKIT channel is shown in D’’’. (E) Correlation plots of MSL3 (green) and DAPI (blue) showing positive correlation between MSL3 and DNA in SSCs (top, gray arrow, ρ=0.76), SPCs (middle, orange arrow, ρ=0.60), and spermatozoa (bottom, yellow arrow, ρ=0.76). (F-F’’) SSC stained for MSL3 (green), DAPI (blue), and SYCP3 (red)shows that meiotic SSC have disparate SYCP3 and MSL3 staining. MSL3 channel is shown in F’ and SYCP3 channel is shown in F’’. (G) A schematic showing that Set2, MSL3, and ATAC complex regulate meiotic progression by transcriptionally regulating synaptonemal complex (SC) components and *ribosomal protein S19b* paralog (*RpS19b*). SC components promote recombination during meiosis and sufficient RpS19 levels is required for Rbfox1 translation which promotes meiotic cell cycle. Together this transcriptional axis promotes the meiotic progression in female *Drosophila*. Scale bar for all images is 20 μm.

## Discussion

Model organisms such as *Drosophila* have given us tremendous insight into how meiosis is regulated, having identified both intrinsic regulators such as translational control factors, as well as extrinsic regulators such as ecdysone signaling that govern this process (Carreira-Rosario et al., 2016; Hongay and Orr-Weaver, 2011; Hughes et al., 2018; Morris and Spradling, 2012; Tastan et al., 2010). However, the transcriptional regulators of the meiotic program in *Drosophila* had yet to be identified. In addition, while meiosis is itself an extremely conserved process, no conserved transcriptional regulator had been identified across organisms (Kimble, 2011). Thus, the overarching questions that needed to be addressed were: 1) What are the transcriptional regulators of meiosis in *Drosophila*? And, 2) Is there a conserved gene regulatory network that controls entry into meiosis?

### The Set2, MSL3, and ATAC transcriptional axis licenses entry into meiosis and its function may be conserved in vertebrates

We have identified Set2, MSL3, and ATAC complex as transcriptional regulators of meiotic entry in *Drosophila*. We demonstrate that loss of either Set2, MSL3, or ATAC complex members in the female germline leads to an accumulation of germ cells that initiate differentiation but stall at the crucial transition step prior to meiotic commitment. We find that the Set2-MSL3-ATAC axis regulates oogenesis downstream of the differentiation factor, Bam, but upstream of the meiotic regulator Rbfox1. The Set2-MSL3-ATAC axis regulates meiosis in two ways: 1) it transcriptionally upregulates members of the synaptonemal complex that is critical to recombination and, 2) it promotes transcription of the germline enriched RpS19 paralog, RpS19b. The expression of RpS19b then controls the translation of *Rbfox1*, which is required for exit from the mitotic cell cycle and entry into meiotic cell cycle (Figure 7G). While several components of the synaptonemal complex are regulated at the transcriptional level, components such as C(3)G and Crossover suppressor on 2 of Manheim (C(2)M) are not. The mRNAs of C(3)G and C(2)M are present in measurable amounts in later stages of oogenesis (24 TPM and 85 TPM, respectively, in whole ovaries) whereas mRNAs of Cona, Ord, and Sunn are restricted to early meiotic stages (8 TPM, <1 TPM, and <1 TPM, respectively, in whole ovaries). This suggests that some synaptonemal complex members such as C(2)M and C(3)G may be regulated at the post-transcriptional level. Taken together, the Set2-MSL3-ATAC complex coordinates transcription of several critical factors of recombination machinery and translation of a meiotic cell cycle regulator to promote entry into meiosis (Figure 7G).

How is expression of critical meiotic genes licensed for expression only during meiosis? We observe that, in the germline, MSL3 expression is restricted to the mitotic and early meiotic stages of oogenesis. We hypothesize that when MSL3 is expressed it functions by binding to Set2 mediated H3K36me3 mark and then recruits a basal transcriptional machinery, ATAC, to enhance transcription of a subset of meiotic genes. Thus, our data suggest that restricted expression of a reader, MSL3, licenses the expression of critical meiotic genes. We do not know what controls expression of MSL3 itself during the mitotic and early meiotic stages. *msl3* mRNA is present as part of the maternal contribution in the egg (Eichhorn et al., 2016; Hua et al., 2014). This suggests that *msl3* mRNA is transcribed in the later stages of oogenesis and is likely post-transcriptionally regulated. While our data demonstrates that MSL3 expression is required for meiotic progression in female *Drosophila*, we do not think MSL3 expression is sufficient for entry into meiosis as overexpression of *msl3* does not lead to precocious meiotic commitment (Figure 2L-M). In addition, H3K36me3 marks are present on gene bodies of transcribed genes, MSL3 is expressed in somatic cells, and ATAC complex is also a basal transcriptional machinery, yet meiotic genes are not expressed in somatic cells (C Santos and Lehmann, 2004; Cinalli et al., 2008; Keogh et al., 2005; Larschan et al., 2007; Marlow, 2015; Morris et al., 2005; Nikolic et al., 2016; Spedale et al., 2012). We predict that a yet unknown factor that is present in the early stages of oogenesis acts in concert with MSL3 to promote expression of meiotic genes. It has been shown that somatic steroid signaling mediated by ecdysone is required for meiotic entry in *Drosophila* (Morris and Spradling, 2012). Indeed, several ecdysone-responsive nuclear receptors are expressed in the germline and are required for its proper development (Belles and Piulachs, 2014, 2015; Carney and Bender, 2000; Schwedes et al., 2011). We speculate that Set2, MSL3, and ATAC could act in concert with ecdysone responsive factor(s) in the germline, that have yet to be identified, to promote entry into meiosis.

MSL3 function in meiotic entry is likely conserved. In mammalian spermatogenesis, STRA8 acts downstream of steroid signaling mediated by RA to promote entry into meiosis (Anderson et al., 2008; Endo et al., 2015; Griswold et al., 2012; Koubova et al., 2006; Zhou et al., 2008). While STRA8 is required in pre-meiotic mammalian spermatogonia to trigger meiosis, its expression outside of its required developmental stage fails to trigger meiosis suggesting it is not sufficient (Kojima et al., 2019). In addition, STRA8 is expressed in Type A spermatogonia to initiate meiosis but it is not clear how the expression of these meiotic genes is sustained. Kojima *et al* have suggested that STRA8 could work in concert with chromatin modifiers such as HATs to promote meiotic gene expression, but such chromatin modifiers have not been identified (Kojima et al., 2019). We find that STRA8 positive spermatocytes express MSL3 (Anderson et al., 2008; Endo et al., 2015; Oulad-Abdelghani et al., 1996; Zhou et al., 2008). This nuclear expression is then maintained for the rest of meiosis (Figure 7A-F’). In addition, MSL3 in mammals is a member of the MSL complex that contains a HAT analogous to the HAT in ATAC complex (Hilfiker et al., 1997; Ravens et al., 2014). Based on MSL3 expression during mouse spermatogenesis, and its function in *Drosophila* oogenesis, we propose that MSL3 could work downstream of steroid signaling from the soma to promote meiotic entry from *Drosophila* to mammals.

### Post-transcriptional control of meiotic commitment

We find MSL3 not only promotes transcription of components of the SC but also promotes entry into meiosis by regulating levels of RpS19 which in turn regulates translation of *Rbfox1*. Rbfox1 then promotes entry into meiosis by repressing mitotic cell cycle and promoting meiotic cell cycle. In mouse, STRA8 also regulates proteins required for meiosis such as synaptonemal complex components as well as post-transcriptional mRNA regulators Meiosis Specific With Coiled-Coil Domain (MEIOC) and YTH Domain-Containing 2 (YTHDC2) (Kojima et al., 2019). MEIOC and YTHDC2 in turn promote entry into meiotic cell cycle (Bailey et al., 2017; Jain et al., 2018; Soh et al., 2017). Thus, coordinated regulation of transcription and translation promotes entry into meiosis in both *Drosophila* and mice, which we propose could be a shared mechanism to modulate entry into meiosis in other organisms.

The germline expresses several unique ribosomal protein paralogs including RpS19b (Gerst, 2018; Marygold et al., 2007). While in other developmental contexts it has been shown that paralogs can play a unique role in translation of specific mRNAs (Desai et al., 2017; Genuth and Barna, 2018a; Herrmann et al., 2013; Segev and Gerst, 2018; Xue and Barna, 2012), our data show that addition of either RpS19b, RpS19a, or hRpS19 can rescue loss of MSL3 phenotype. This suggests that *Rbfox1* translation, which promotes meiotic entry, is particularly sensitive to levels of RpS19 but not specific paralogs (Carreira-Rosario et al., 2016; Tastan et al., 2010). While we find that RpS19 a and b are incorporated into the ribosome and regulates translation of *Rbfox1,* we cannot exclude the possibility that RpS19 regulates translation of *Rbfox1* via an extra-ribosomal function. Thus, expression of MSL3 causes an upregulation of ribosomal protein S19 to promote entry into meiosis.

Levels of ribosomal proteins, including RpS19, affecting translation of specific transcripts has precedence in mammals and *Drosophila* (Genuth and Barna, 2018b; Khajuria et al., 2018; Kondrashov et al., 2011; Kong et al., 2019; Palumbo et al., 2017; Segev and Gerst, 2018; Shi et al., 2017; Signer et al., 2014; Simsek et al., 2017; Xue et al., 2015; Xue and Barna, 2012). Mutations in RpS19 result in ribosomopathies including Diamond-Blackfan anemia (DBA). It has been shown that in DBA, due to loss of RpS19, results in loss of translation of GATA1 transcription factor which causes a failure of hematopoietic stem cell differentiation (Draptchinskaia et al., 1999; Gazda et al., 2004; Khajuria et al., 2018; Ludwig et al., 2014; Willig et al., 2000). In addition, during mouse development, *Ribosomal protein L38* (*RpL38*) is expressed at higher levels in specific tissues such as developing vertebrae. Reduction in levels of *RpL38* in mice results in decrease of *hox* mRNA translation leading to homeotic transformations of the vertebrae (Kondrashov et al., 2011). Intriguingly, *RpL38* and *RpS19* are among the many ribosomal proteins that are differentially expressed in various tissues (Marygold et al., 2007; Xue and Barna, 2012). Thus, not only can ribosomal protein paralogs affect translation of specific mRNAs, but levels of particular ribosomal proteins can also alter translation of specific transcripts which in turn dictates developmental outcomes by regulating cell fate (Khajuria et al., 2018; Kondrashov et al., 2011). Our work outlines a mechanism by which levels of specific ribosomal proteins can be developmentally regulated to control gene expression programs.

## Acknowledgments

We are grateful to all members of the Rangan laboratory for discussion and comments on the manuscript. Additionally, we would like to thank Dr. Mitzi Kuroda, Dr. Erika Bach, Dr. Ruth Lehmann, Dr. Scott Hawley, Dr. Sharon Bickel, Dr. Gaby Fuchs, Dr. Melinda Larsen, Bloomington Drosophila Stock Center, Vienna Drosophila Resource Center, Transgenic RNAi Project (NIH/NIGMS R01-GM084947), The BDGP Gene Disruption Project, and FlyBase for fly stocks and reagents. Furthermore, we would like to thank CFG Facility at the University at Albany (UAlbany) for performing RNA-seq and polysome-seq analyses. Lastly, we are thankful for both Dr. Marlene Belfort and Dr. Gaby Fuchs for their generosity in allowing us to use instruments. P.R. acknowledges funding from NIH/NIGMS 2R01GM111770–06 and M.B. acknowledges funding from R01 GM125812. P.F. is supported by NICHD and NIH under the award numbers 1R15HD09641101 and 1R01HD097331-01 (PF) and by the NIDCD of NIH under award number 1R01DC017149-01A1.

## Contributions

A.M., K.S., and P.R. designed experiments, analyzed, and interpreted data. E.T.M. provided bioinformatic support. A.M., K.S., and M.U. performed *Drosophila* experiments; J.J., J.M.L., and P.F. designed and performed mouse experiments. N.D.M., S.J., and M.B. made RpS19a and RpS19b fly lines. A.M. and P.R. wrote the manuscript, which all authors edited and approved.

## Materials and Methods

### Fly lines

Flies were grown at 25-31°C and dissected between 1-5 days post-eclosion.

The following RNAi stocks were used in this study; if more than one line is listed, then both were quantitated and the first was shown in the main figure: *Set2* RNAi (Bloomington #33706 and #42511), *msl3* RNAi (Bloomington #35272), *NC2β* RNAi (Bloomington #57421 and VDRC #v3161), *Ada2a* RNAi (Bloomington #50905), *Atac1* RNAi (VDRC #v36092), *Atac2* RNAi (VDRC #v16047), *D12* RNAi (VDRC #v29954), *wds* RNAi (Bloomington #60399), *NC2α* RNAi (Bloomington #67277), *bam* RNAi (Bloomington #58178), *hs-bam*/TM3 (Bloomington #24637) *RpS19b* RNAi (VDRC #v22073 and #v102171), and *RpS19a* RNAi (Bloomington #42774 and VDRC #v107188).

The following mutant and overexpression stocks were used in this study: *Set2^1^*/FM7 (Bloomington #77916), *msl3^1^*/TM3 (Bloomington #5872), *msl3^KG^*/TM3 (Bloomington #13165), *mls3^MB^*/TM3 (Bloomington #29244), *msl1^γ216^*/CyO (Bloomington #5870), *msl1^kmB^*/CyO (Bloomington #25157), *msl2^227^*/CyO (Bloomington #5871), *msl2^kmA^*/CyO (Bloomington #25158), *mle^1^*/SM1 (Bloomington #4235), *mle^9^*/CyO (Bloomington #5873), *Hel89B^08724^*/TM3 (Bloomington #11732), *Hel89B Df*/TM6 (Bloomington #7982), *Atac2^e03046^*/CyO (Bloomington #18111), *RpS19b^EY00801^* (Bloomington #15043), *RpS19b^CRISPR^* (this study), *UAS-hRpS19-HA* (Bloomington #66014), and *UAS-msl3-GFP* (this study).

The following tagged lines were used in this study: *msl3-GFP* (Kuroda Lab), *RpS19a-3xHA* (this study), *RpS19b-GFP* (this study), and *ord-GFP* (Bickel Lab).

The following tissue-specific drivers were used in this study: *UAS-Dcr2;nosGAL4* (Bloomington #25751), *UAS-Dcr2;nosGAL4;bam-GFP* (Lehmann Lab), *nosGAL4;MKRS*/TM6 (Bloomington #4442), and *If*/CyO*;nosGAL4* (Lehmann Lab).

### Dissection and Immunostaining

Ovaries were dissected and stained as previously described (McCarthy et al., 2018b). The following primary antibodies were used: mouse anti-1B1 (1:20; DSHB), Rabbit anti-Vasa (1:1,000; Rangan Lab), Chicken anti-Vasa (1:1,000 (Upadhyay et al., 2016)), Rabbit anti-GFP (1:2,000; abcam, ab6556), Guinea pig anti-Rbfox1 (1:1,000 (Tastan et al., 2010)), Mouse anti-C(3)G (1:1000; Hawley Lab), Rabbit anti-H3K36me3 (1:500; abcam, ab9050), Rabbit anti-pMAD (1:150; abcam, ab52903), Mouse anti-BamC (1:200; DSHB, Supernatant), Rabbit anti-Bru (1:500; Lehmann Lab), Rabbit anti-Egl (1:1,000; Lehmann Lab), Rat anti-HA (1:500; Roche, 11 867 423 001), and Rabbit anti-RpS19 (1:20; Proteintech, 15085-1-AP). Anti-RpS19 was pre-cleared at 1:20, the supernatant was then diluted at 1:2.5 for staining. The following secondary antibodies were used: Alexa 488 (Molecular Probes), Cy3 and Cy5 (Jackson Labs) were used at a dilution of 1:500.

### Fluorescence Imaging

The tissues were visualized, and images were acquired using a Zeiss LSM-710 confocal microscope under 20X, 40X and 63X oil objective.

### AU quantification of protein or *in situ*

To quantify antibody staining intensities for Rbfox1, H3K36me3, Bruno, GFP, HA, and RpS19 or *in situ* probe fluorescence in germ cells, images for both control and experimental germaria were taken using the same confocal settings. Z stacks were obtained for all images. Similar planes in control and experimental germaria were chosen, the area of germ cells positive for the proteins or *in situs* of interest was outlined and analyzed using the ‘analyze’ tool in Fiji (ImageJ). The mean intensity and area of the specified region was obtained. An average of all the ratios (Mean/Area), for the proteins or *in situs* of interest, per image was calculated for both, control and experimental. Germline intensities were normalized to somatic intensities or if the protein or *in situ* of interest is germline enriched and not expressed in the soma they were normalized to Vasa or background. The highest mean intensity between control and experimental(s) was used to normalize to a value of 1 A.U. on the graph. A minimum of 5 germaria was used for quantitation.

### Egg laying assays

Assays were conducted in cages with females under testing and wild type control males. Cages were maintained at 25°C. All flies were 1 day post-eclosion upon setting up the experiment and analyses were performed on four consecutive days. The number of eggs laid were normalized to the total number of females.

### RNA-seq library preparation and analysis

Ovaries from flies were dissected in 1x PBS. RNA was isolated using TRIzol (Invitrogen, 15596026), treated with DNase (TURBO DNA-free Kit, Life Technologies, AM1907), and then run on a 1% agarose gel to check integrity of the RNA. To generate mRNA enriched libraries, total RNA was treated with poly(A)tail selection beads (Bioo Scientific Corp., NOVA-512991) and then following the manufacturer’s instructions of the NEXTflex Rapid Directional RNA-seq Kit (Bioo Scientific Corp., NOVA-5138-08), except that RNA was fragmented for 13 min. Single-end mRNA sequencing (75 base pair) was performed on biological duplicates from each genotype on an Illumina NextSeq500 by the Center for Functional Genomics (CFG).

After quality assessment, the sequenced reads were aligned to the *Drosophila melanogaster* genome (UCSCdm6) using HISAT2 (version 2.1.0) with the RefSeq-annotated transcripts as a guide (Kim et al., 2015). Raw counts were generated using featureCounts (version 1.6.0.4) (Liao et al., 2014). Differential gene expression was assayed by edgeR (version 3.16.5), using a false discovery rate (FDR) of 0.05, and genes with fourfold or higher were considered significant. The raw and unprocessed data for RNA-seq generated during this study are available at Gene Expression Omnibus (GEO) databank under accession number: XXXXX.

### *In situ* hybridization

Adult ovaries (5 ovary pairs per sample per experiment) were dissected and fixed as previously described. The ovaries were washed with PT (1x phosphate-buffered saline (PBS), 0.1% Triton-X 100) 3 times for 5 minutes each. Ovaries were permeabilized by washing once with increasing concentrations of methanol for 5 minutes each (30% methanol in PT, 50% methanol in PT, and 70% methanol in PT) then incubating in methanol for 10 minutes. Ovaries were then post-fixed by washing once with decreasing concentrations of methanol for 5 minutes each (70% methanol in PT, 50% methanol in PT, and 30% methanol in PT). Ovaries were then washed with PT 3 times for 5 minutes and then pre-hybridized in wash buffer for 10 minutes (10% deionized formamide and 10% 20x SSC in RNase-free water). Ovaries were incubated overnight in hybridization solution (10% dextran sulfate, 1 mg/ml yeast tRNA, 2 mM RNaseOUT, 0.02 mg/ml BSA, 5x SSC, 10% deionized formamide, and RNase-free water) at 30°C. The hybridization solution was removed, and ovaries washed with Wash Buffer 2 times for 30 minutes at 30°C. Wash Buffer was removed, and ovaries were mounted using Vectashield with 4’,6’-diamidino-2-phenylindole (DAPI).

### *In situ* probe design and generation

Templates were amplified with gene specific primers (listed below) and then followed manufacturer’s instructions of Thermo Fisher’s FISH tag RNA kit (F32954) for generating fluorescently labeled probes.

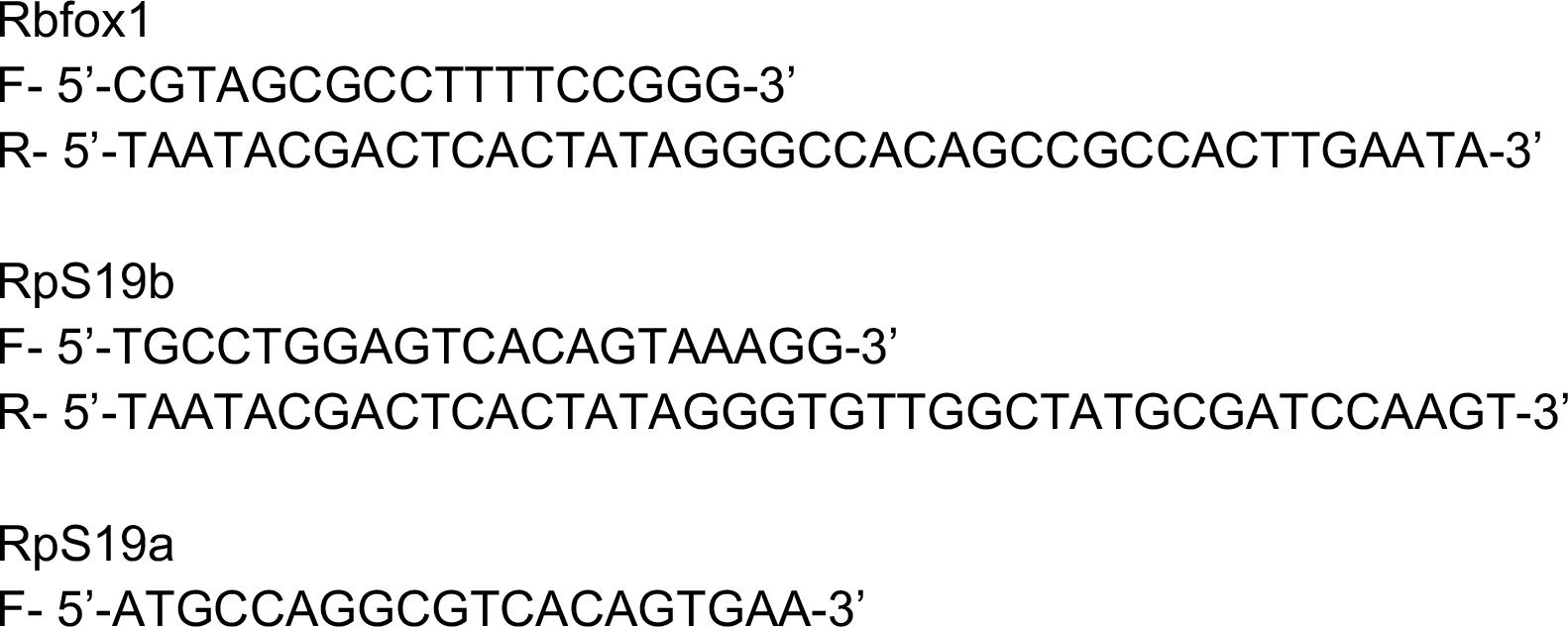

### Measurement of global protein synthesis

Protein synthesis was detected using short-term ovary incorporation assay, Click-iT Plus OPP (Invitrogen, C10456). Ovaries were dissected in Schneider’s *Drosophila* media (Thermo Fisher, 21720024) and then incubated in 50 μM OPP reagent for 30 minutes. Tissue was washed in 1x PBS and then fixed for 15 min in 1x PBS plus 5% methanol-free formaldehyde. Tissue was then permeabilized with 1% Triton X-100 in 1x PBST (1x PBS with 0.2% Tween 20) for 30 minutes, samples were then washed in 1x PBS and were incubated in Click-iT reaction cocktail following the manufacturer’s instructions. Samples were washed with Click-iT reaction rise buffer and then immunostained following previously described procedures.

### Generating fly lines

#### CRISPR mutant

To generate the *RpS19b* mutants, guide RNAs were designed using http://tools.flycrispr.molbio.wisc.edu/targetFinder and synthesized as 5-unphosphorylated oligonucleotides, annealed, phosphorylated, and ligated into the BbsI sites of the pU6-BbsI-chiRNA vector using the primers listed below (Gratz et al., 2013). Homology arms were synthesized as a gene block (IDTDNA) and cloned into pHD-dsRed-attP ((Gratz et al., 2015); Addgene) using Gibson Assembly (gene blocks listed in Supplementary Methods). Guide RNAs and the donor vector were co-injected into *nos-Cas9* embryos (Rainbow Transgenics).

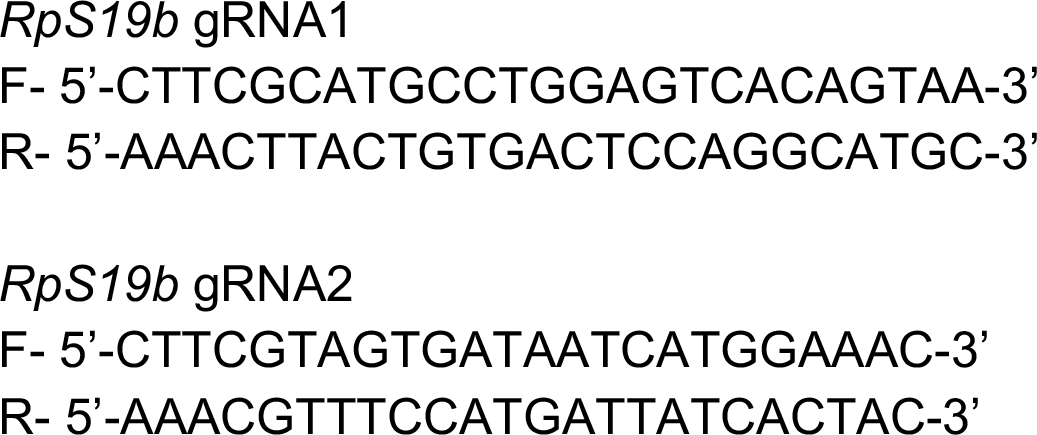

#### *RpS19a-3xHA* and *RpS19b-GFP* tagged lines

*RpS19a3x-HA* (referred to as *RpS19a-HA* throughout text) and *RpS19b-GFP* tagged lines were made using a combination of *in vivo* bacterial recombineering and GatewayTM Technology as previously described (Shalaby et al., 2017).

#### *UAS-msl3-GFP* overexpression line

RNA was extracted from *w^1118^* ovaries and made into cDNA using a SuperScript II-Strand Kit (Thermo Fisher, 18064014). *msl3* CDS was amplified, *att*B sites and tagged sequence was amplified into the PCR product using the primers listed below. PCR products were cloned into pDONR (Thermo Fisher, 11789-020) and swapped into pENTR (Thermo Fisher, 11791-020) using BP and LR reactions, respectively. The plasmid was sent for injection into *w^1118^* flies (Genetic Services).

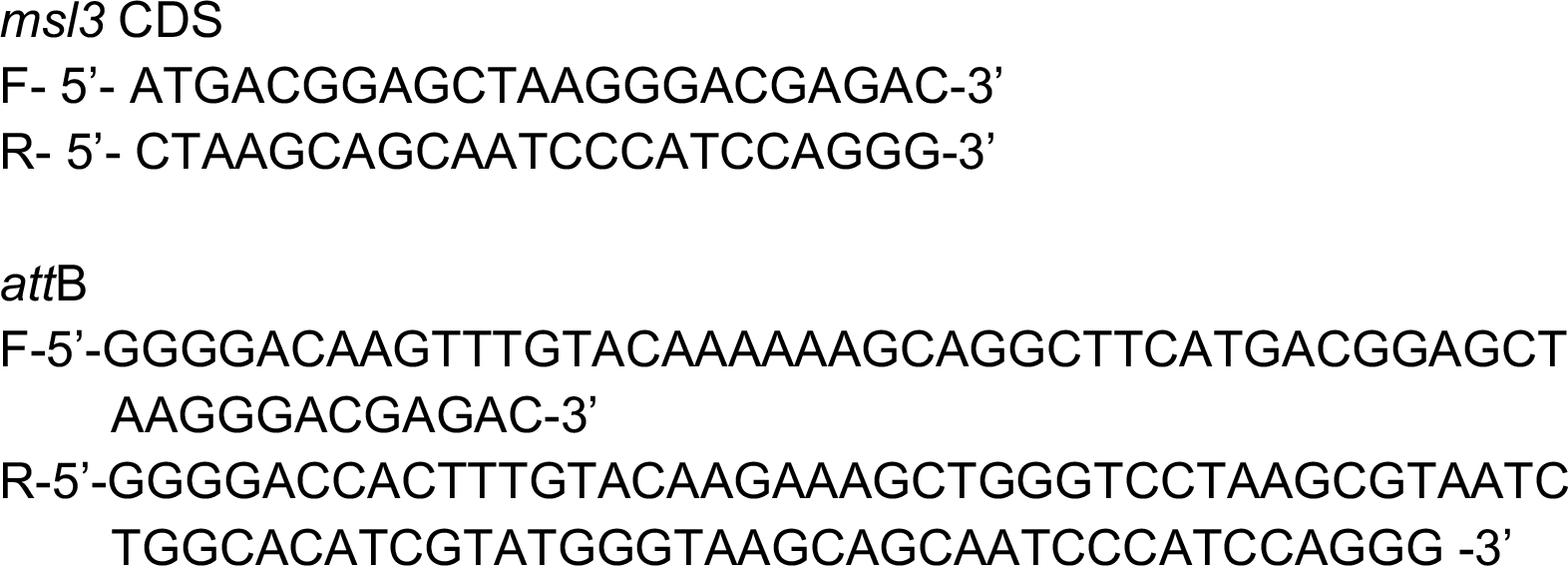

### Polysome profiling and polysome-seq

Polysome profiling of ovaries from *bam* RNAi*;hs-bam* and *UAS-Dcr2;nosGAL4 >RpS19b* RNAi flies was adapted from (Flora et al., 2018; Fuchs et al., 2011). 200 ovary pairs were dissected in Schneider’s media and immediately flash frozen with liquid nitrogen. Ovaries were homogenized in Lysis Buffer, 20% of lysate was used as input for mRNA isolation and library preparation (as described above). Samples were loaded onto 10-50% CHX-supplemented sucrose gradients in 9/16 x 3.5 PA tubes (Beckman Coulter, #331372) and spun at 35,000 x g in SW41 for 2.45-3 hours at 4°C. Gradients were fractionated with a Density Gradient Fractionation System (#621140007). RNA was extracted using acid phenol-chloroform and precipitated overnight. Pelleted RNA was resuspended in 20 μL water and libraries were prepared as described above.

### Western blot

50-200 CB enriched *RpS19a-HA* and *RpS19b-GFP* ovaries and 30 adult *RpS19b-GFP;RpS19a-HA* ovaries were dissected and prepared as described above except sucrose solutions were supplemented with either 100 μg/μL CHX or 2 mM puromycin with 1 mg heparin prior to making gradients. Following fractionation, protein was extracted by ethanol precipitation and run on a TGX pre-cast gradient gel (BioRad, #456-1094). Blots were blocked with 5% milk in 1x PBST and incubated in primary antibody in 5% BSA in 1x PBST. Following 1x PBST washing, blots were incubated in secondary antibody in 5% milk in 1x PBST. Blots were washed with 1x PBST and then imaged with chemi-luminescence kit (BioRad, #170-5060). The following primary antibodies were used: Rabbit anti-GFP (1:4,000; abcam, ab6556), Rat anti-HA (1:3,000; Roche, 11 867 423 001), Rabbit anti-RpS25 (1:1,000; abcam, ab40820), and Rabbit anti-RpS19 (1:1,000; Proteintech, 15085-1-AP). The following secondary antibodies were used: anti-Rat HRP (1:10,000; Jackson Labs, 112-035-003) and anti-Rabbit HRP (1:10,000; Jackson Labs, 111-035-144).

### Statistical Analysis

Relative fluorescence signals were compared between control and experimental groups using parametric tests (Student t-test or one-way ANOVA). Horizontal lines on scatter dot plots represent mean with 95% confidence interval and stars on stacked bar graphs represent statistical significance of corresponding color data set. Reported p-values correspond to two-tailed tests. Analysis of percentage defect were compared between control and experimental groups using Fisher’s exact test. All analyses were performed using Prism 8 software (GraphPad) and reported in figure legends.

### Materials and reagents for fly husbandry

Fly food was made by using previously described procedures (Upadhyay et al., 2018).

### Mice

#### Ethics statement

Collection and use of mouse specimens for this study were approved by the Institutional Animal Care and Use Committee (IACUC) at The University at Albany.

The mice used in this study were adult males on a CD-1 background. All data were collected from mice kept under similar housing conditions, in transparent cages on a normal 12 hr. light/dark cycle.

#### Dissection and tissue preparation of mouse testes

Tissue was collected from adult male mice on a CD-1 background (Forni, 2006). The mice were perfused first with 1x PBS then with 3.7% formaldehyde in 1x PBS. Testes were isolated at the time of perfusion and immersion-fixed for 2-3 hours at 4°C. The samples were then cryoprotected in 30% sucrose in 1x PBS overnight at 4°C then embedded in Tissue-Tek O.C.T. Compound (Sakura Finetek, 4583) using dry ice, and stored at −80°C.

Tissue was cryosectioned (Leica Cryostat, CM3050S) at 20 μm and collected on microscope slides (VWR, 48311-703) for immunostainings. All slides were stored at - 80°C until ready for staining.

#### Immunostaining of mouse testes

Citrate buffer (pH 6.0) antigen retrieval was performed before immunostaining. Tissue was incubated in blocking solution (10% horse serum, 1% BSA, 0.5% Triton X-100, and 0.1% Sodium Azide) for 40 minutes up until 1 hour at room temperature. The following primary antibodies were used: Rabbit anti-MSL3 (1:500; Invitrogen, PA5-56967), Goat anti-cKIT (1:250; R&D Systems, AF1356), Rabbit anti-Stra8 (1:250; abcam, ab49602), and Mouse anti-SYCP3 (1:500; abcam, ab97672).

The following secondary antibodies were used: Alexa Fluor 488, Alexa Fluor 594, Alexa Fluor 680 were at a dilution of 1:1000 (Molecular Probes and Jackson ImmunoResearch Laboratories). Sections were counterstained with DAPI (1:3000; Sigma-Aldrich, 28718-90-3) and coverslips were mounted with FluoroGel (Electron Microscopy Services, 17985-10). Confocal microscopy pictures were taken on a Zeiss LSM 710 microscope using a 40x oil objective.

All unique/stable reagents generated in this study are available from the Lead Contact without restriction.

