## Supplemental Files for "MSL3 coordinates a transcriptional and translational meiotic program in female Drosophila"

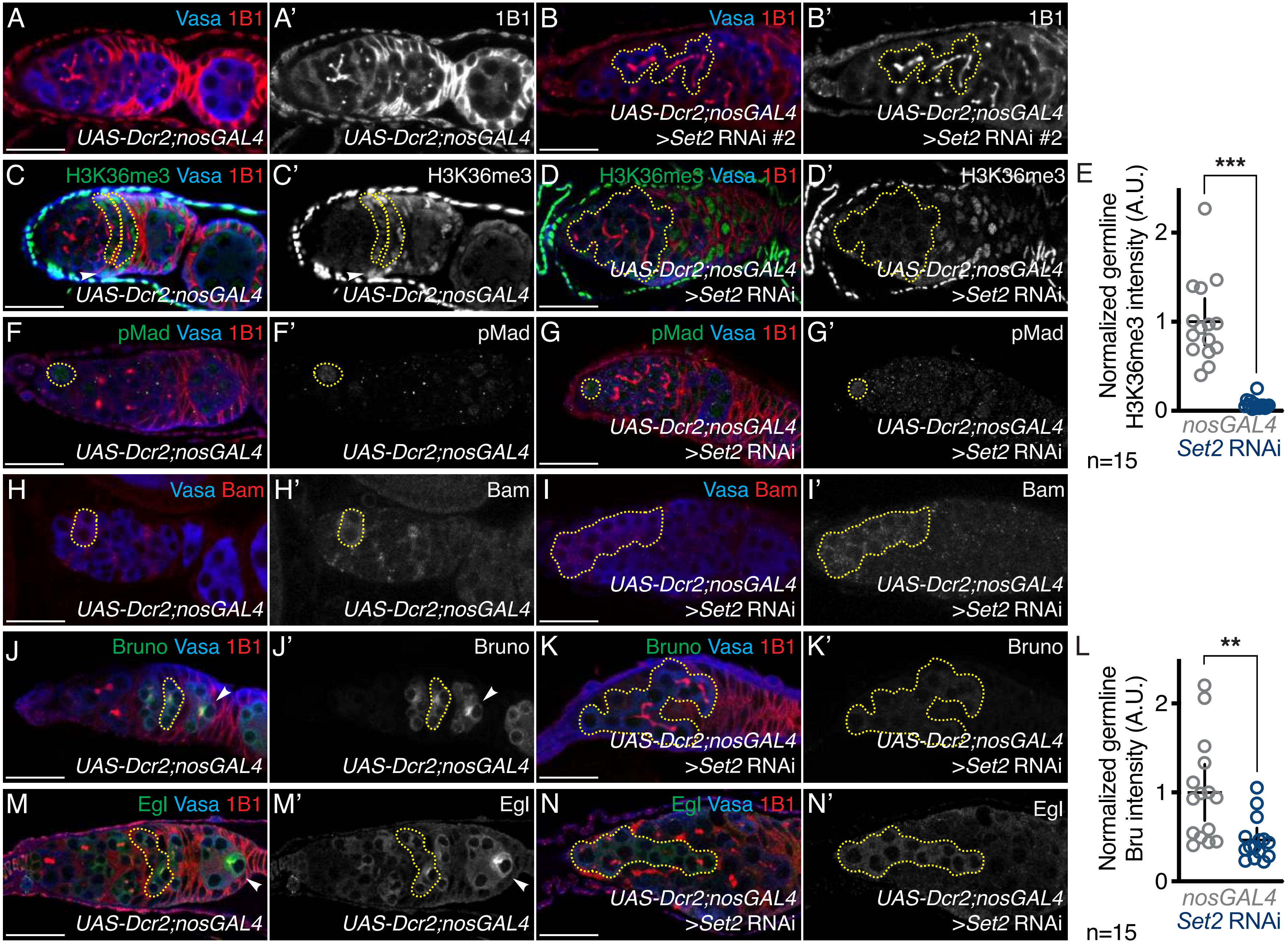
A
Vasa 1B1
UAS-Dcr2;nosGAL4
A'
1B1
UAS-Dcr2;nosGAL4
B
Vasa 1B1
UAS-Dcr2;nosGAL4 >Set2 RNAi #2
B'
1B1
UAS-Dcr2;nosGAL4 >Set2 RNAi #2
C
H3K36me3 Vasa 1B1
UAS-Dcr2;nosGAL4
C'
H3K36me3
UAS-Dcr2;nosGAL4
D
H3K36me3 Vasa 1B1
UAS-Dcr2;nosGAL4 >Set2 RNAi
D'
H3K36me3
UAS-Dcr2;nosGAL4 >Set2 RNAi
E
***
Normalized germline H3K36me3 intensity (A.U.)
0
1
2
nosGAL4
Set2 RNAi
n=15
F
pMad Vasa 1B1
UAS-Dcr2;nosGAL4
F'
pMad
UAS-Dcr2;nosGAL4
G
pMad Vasa 1B1
UAS-Dcr2;nosGAL4 >Set2 RNAi
G'
pMad
UAS-Dcr2;nosGAL4 >Set2 RNAi
H
Vasa Bam
UAS-Dcr2;nosGAL4
H'
Bam
UAS-Dcr2;nosGAL4
I
Vasa Bam
UAS-Dcr2;nosGAL4 >Set2 RNAi
I'
Bam
UAS-Dcr2;nosGAL4 >Set2 RNAi
J
Bruno Vasa 1B1
UAS-Dcr2;nosGAL4
J'
Bruno
UAS-Dcr2;nosGAL4
K
Bruno Vasa 1B1
UAS-Dcr2;nosGAL4 >Set2 RNAi
K'
Bruno
UAS-Dcr2;nosGAL4 >Set2 RNAi
L
**
Normalized germline Bru intensity (A.U.)
0
1
2
nosGAL4
Set2 RNAi
n=15
M
Egl Vasa 1B1
UAS-Dcr2;nosGAL4
M'
Egl
UAS-Dcr2;nosGAL4
N
Egl Vasa 1B1
UAS-Dcr2;nosGAL4 >Set2 RNAi
N'
Egl
UAS-Dcr2;nosGAL4 >Set2 RNAi

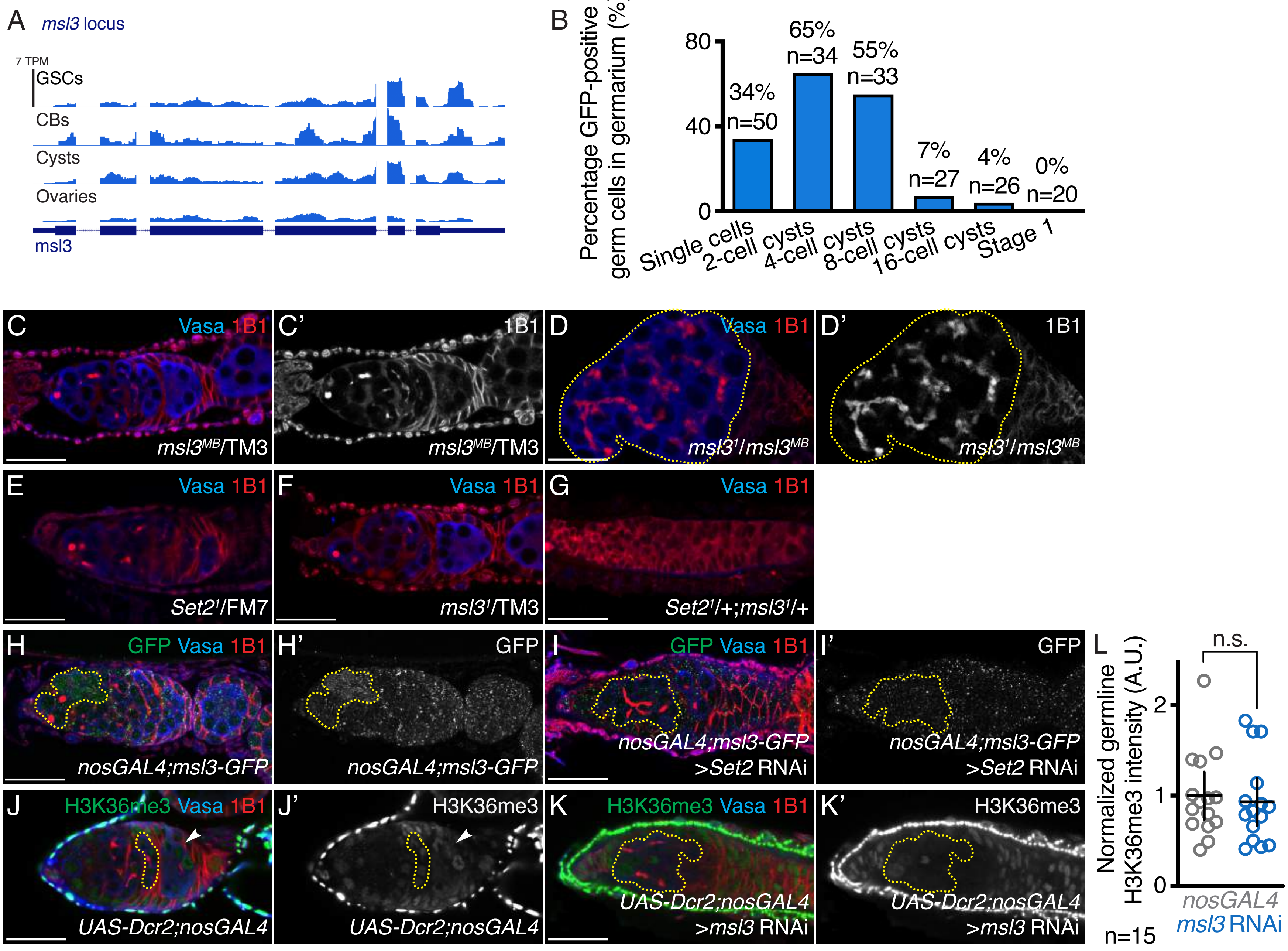
A
msl3 locus
7 TPM
GSCs
CBs
Cysts
Ovaries
msl3
B
Percentage GFP-positive germ cells in germarium (%)
80
40
0
34% n=50
65% n=34
55% n=33
7% n=27
4% n=26
0% n=20
Single cells
2-cell cysts
4-cell cysts
8-cell cysts
16-cell cysts
Stage 1
C Vasa 1B1
msl3MB/TM3
C' 1B1
msl3MB/TM3
D Vasa 1B1
msl31/msl3MB
D' 1B1
msl31/msl3MB
E Vasa 1B1
Set21/FM7
F Vasa 1B1
msl31/TM3
G Vasa 1B1
Set21/+;msl31/+
H GFP Vasa 1B1
nosGAL4;msl3-GFP
H' GFP
nosGAL4;msl3-GFP
I GFP Vasa 1B1
nosGAL4;msl3-GFP >Set2 RNAi
I' GFP
nosGAL4;msl3-GFP >Set2 RNAi
J H3K36me3 Vasa 1B1
UAS-Dcr2;nosGAL4
J' H3K36me3
UAS-Dcr2;nosGAL4
K H3K36me3 Vasa 1B1
UAS-Dcr2;nosGAL4 >msl3 RNAi
K' H3K36me3
UAS-Dcr2;nosGAL4 >msl3 RNAi
L
Normalized germline H3K36me3 intensity (A.U.)
n.s.
2
1
0
nosGAL4
msl3 RNAi
n=15

Figure 2-Supplement 2

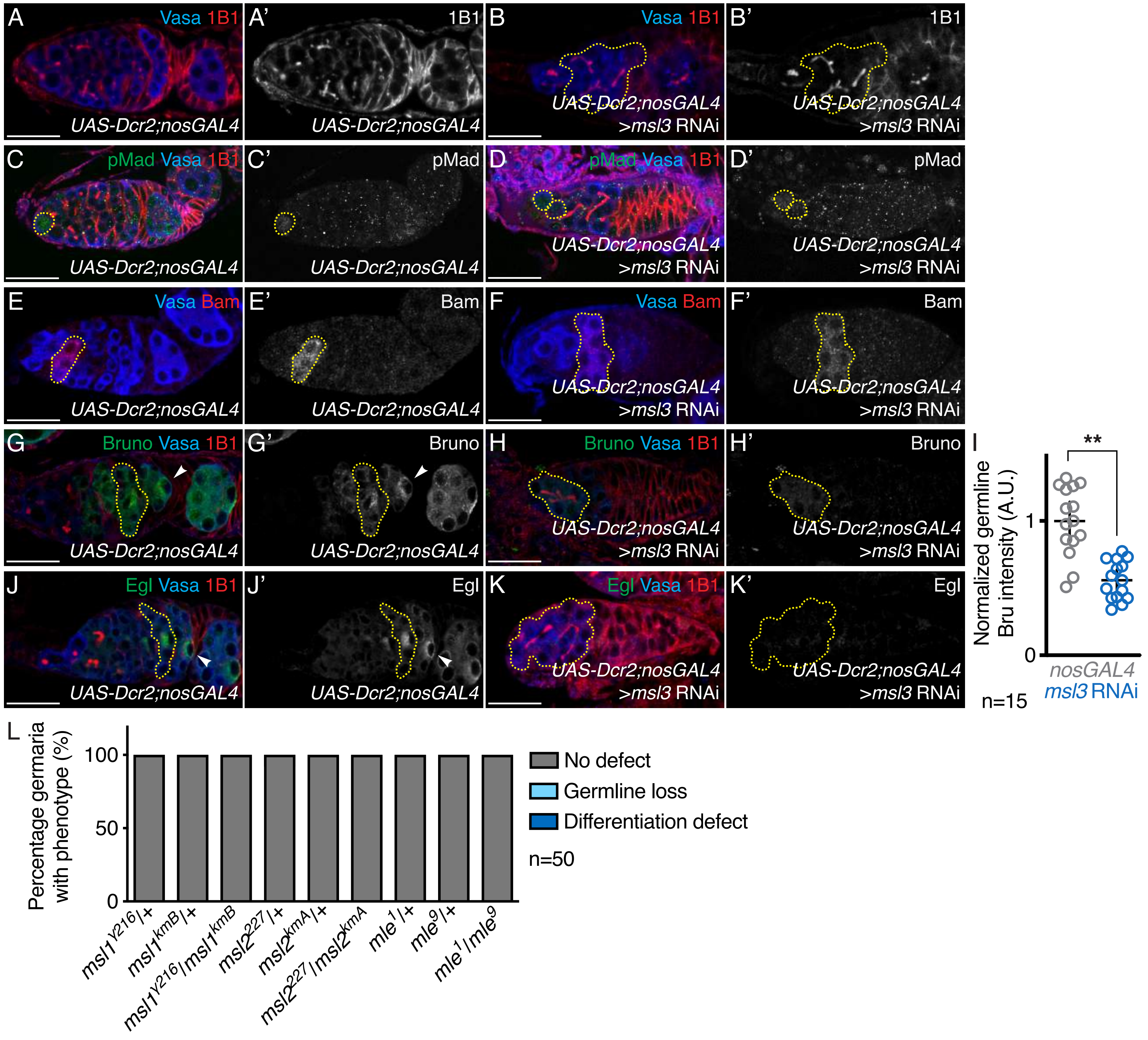

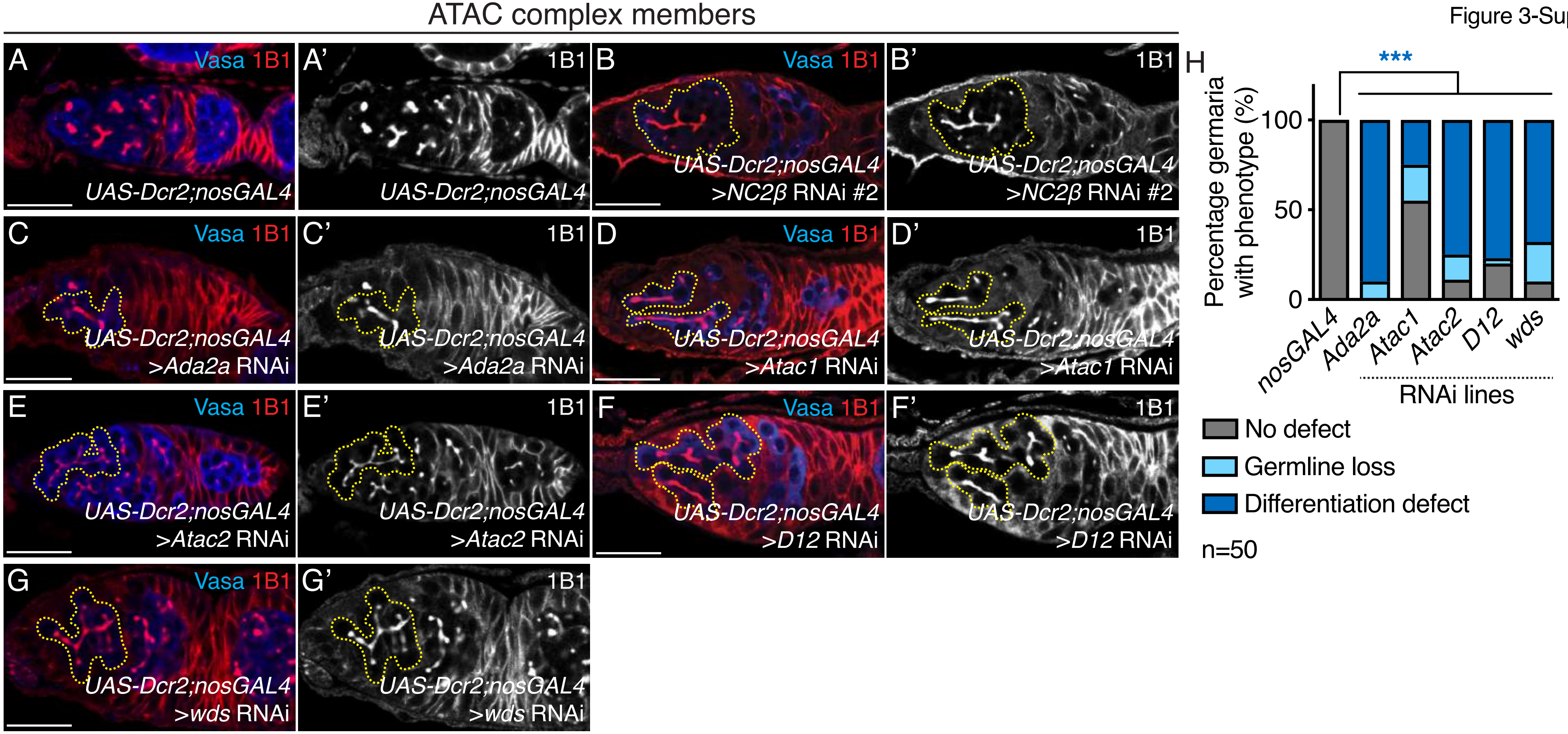
ATAC complex members
A
Vasa 1B1
UAS-Dcr2;nosGAL4
A'
1B1
UAS-Dcr2;nosGAL4
B
Vasa 1B1
UAS-Dcr2;nosGAL4
>NC2β RNAi #2
B'
1B1
UAS-Dcr2;nosGAL4
>NC2β RNAi #2
C
Vasa 1B1
UAS-Dcr2;nosGAL4
>Ada2a RNAi
C'
1B1
UAS-Dcr2;nosGAL4
>Ada2a RNAi
D
Vasa 1B1
UAS-Dcr2;nosGAL4
>Atac1 RNAi
D'
1B1
UAS-Dcr2;nosGAL4
>Atac1 RNAi
E
Vasa 1B1
UAS-Dcr2;nosGAL4
>Atac2 RNAi
E'
1B1
UAS-Dcr2;nosGAL4
>Atac2 RNAi
F
Vasa 1B1
UAS-Dcr2;nosGAL4
>D12 RNAi
F'
1B1
UAS-Dcr2;nosGAL4
>D12 RNAi
G
Vasa 1B1
UAS-Dcr2;nosGAL4
>wds RNAi
G'
1B1
UAS-Dcr2;nosGAL4
>wds RNAi
H
***
Percentage germaria with phenotype (%)
100
50
0
nosGAL4
Ada2a
Atac1
Atac2
D12
wds
RNAi lines
No defect
Germline loss
Differentiation defect
n=50

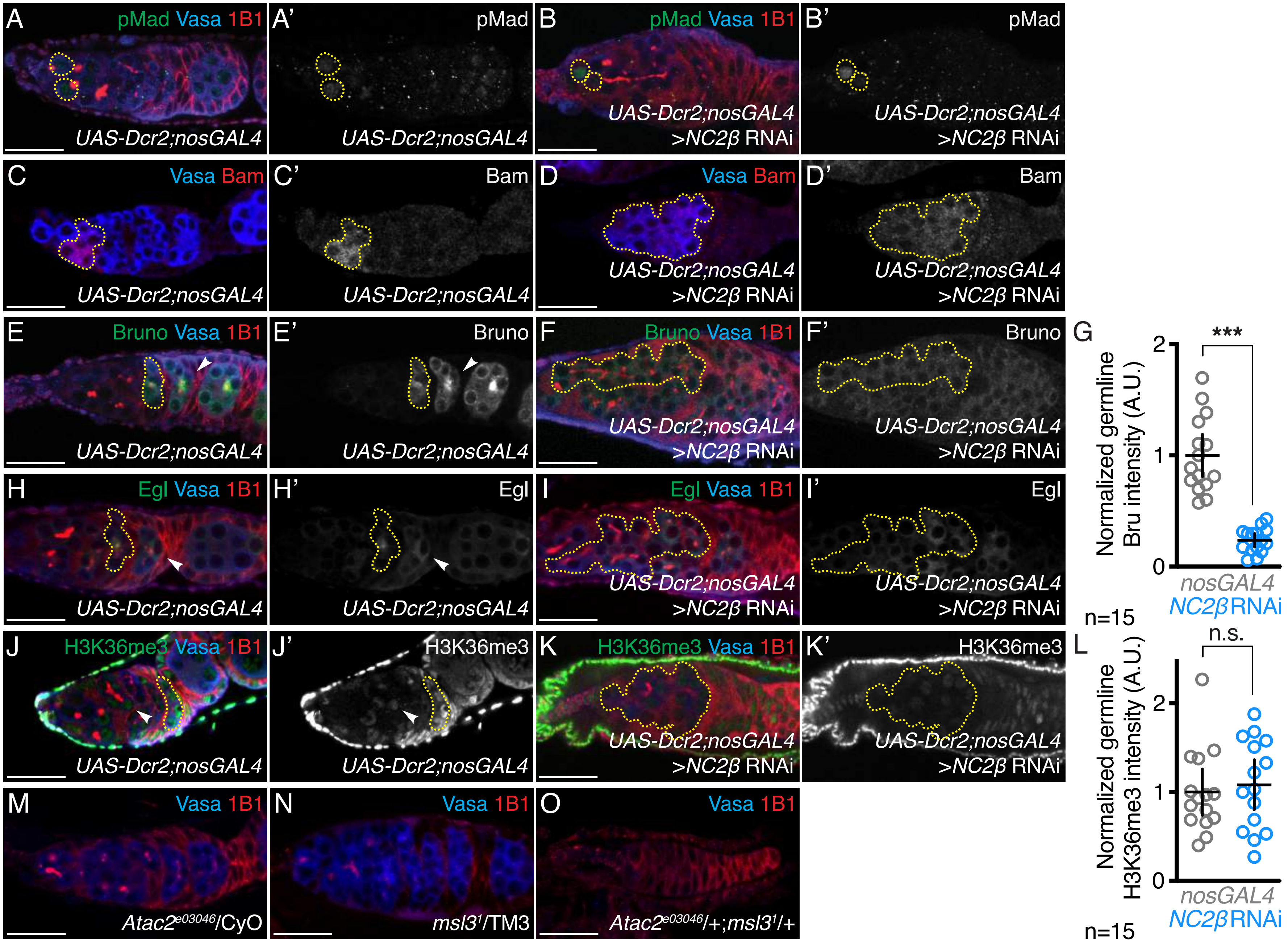
A
pMad Vasa 1B1
UAS-Dcr2;nosGAL4
A'
pMad
UAS-Dcr2;nosGAL4
B
pMad Vasa 1B1
UAS-Dcr2;nosGAL4
>NC2β RNAi
B'
pMad
UAS-Dcr2;nosGAL4
>NC2β RNAi
C
Vasa Bam
UAS-Dcr2;nosGAL4
C'
Bam
UAS-Dcr2;nosGAL4
D
Vasa Bam
UAS-Dcr2;nosGAL4
>NC2β RNAi
D'
Bam
UAS-Dcr2;nosGAL4
>NC2β RNAi
E
Bruno Vasa 1B1
UAS-Dcr2;nosGAL4
E'
Bruno
UAS-Dcr2;nosGAL4
F
Bruno Vasa 1B1
UAS-Dcr2;nosGAL4
>NC2β RNAi
F'
Bruno
UAS-Dcr2;nosGAL4
>NC2β RNAi
G
Normalized germline Bru intensity (A.U.)
***
2
1
0
nosGAL4
NC2β RNAi
n=15
H
Egl Vasa 1B1
UAS-Dcr2;nosGAL4
H'
Egl
UAS-Dcr2;nosGAL4
I
Egl Vasa 1B1
UAS-Dcr2;nosGAL4
>NC2β RNAi
I'
Egl
UAS-Dcr2;nosGAL4
>NC2β RNAi
J
H3K36me3 Vasa 1B1
UAS-Dcr2;nosGAL4
J'
H3K36me3
UAS-Dcr2;nosGAL4
K
H3K36me3 Vasa 1B1
UAS-Dcr2;nosGAL4
>NC2β RNAi
K'
H3K36me3
UAS-Dcr2;nosGAL4
>NC2β RNAi
L
Normalized germline H3K36me3 intensity (A.U.)
n.s.
2
1
0
nosGAL4
NC2β RNAi
n=15
M
Vasa 1B1
Atac2e03046/CyO
N
Vasa 1B1
msl31/TM3
O
Vasa 1B1
Atac2e03046/+;msl31/+

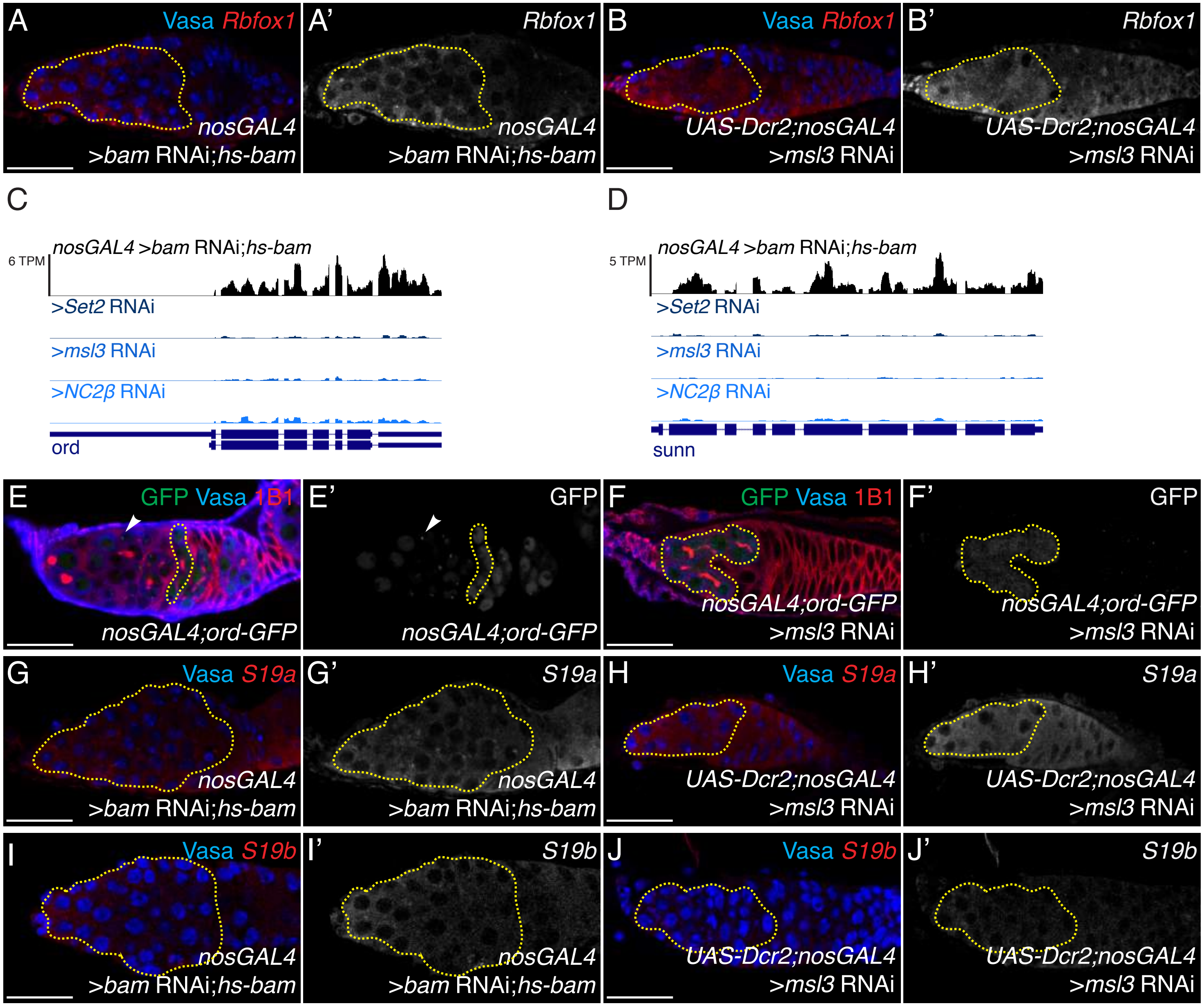
A
Vasa Rbfox1
nosGAL4
>bam RNAi;hs-bam
A'
Rbfox1
nosGAL4
>bam RNAi;hs-bam
B
Vasa Rbfox1
UAS-Dcr2;nosGAL4
>msl3 RNAi
B'
Rbfox1
UAS-Dcr2;nosGAL4
>msl3 RNAi
C
6 TPM
nosGAL4 >bam RNAi;hs-bam
>Set2 RNAi
>msl3 RNAi
>NC2β RNAi
ord
D
5 TPM
nosGAL4 >bam RNAi;hs-bam
>Set2 RNAi
>msl3 RNAi
>NC2β RNAi
sunn
E
GFP Vasa 1B1
nosGAL4;ord-GFP
E'
GFP
nosGAL4;ord-GFP
F
GFP Vasa 1B1
nosGAL4;ord-GFP
>msl3 RNAi
F'
GFP
nosGAL4;ord-GFP
>msl3 RNAi
G
Vasa S19a
nosGAL4
>bam RNAi;hs-bam
G'
S19a
nosGAL4
>bam RNAi;hs-bam
H
Vasa S19a
UAS-Dcr2;nosGAL4
>msl3 RNAi
H'
S19a
UAS-Dcr2;nosGAL4
>msl3 RNAi
I
Vasa S19b
nosGAL4
>bam RNAi;hs-bam
I'
S19b
nosGAL4
>bam RNAi;hs-bam
J
Vasa S19b
UAS-Dcr2;nosGAL4
>msl3 RNAi
J'
S19b
UAS-Dcr2;nosGAL4
>msl3 RNAi

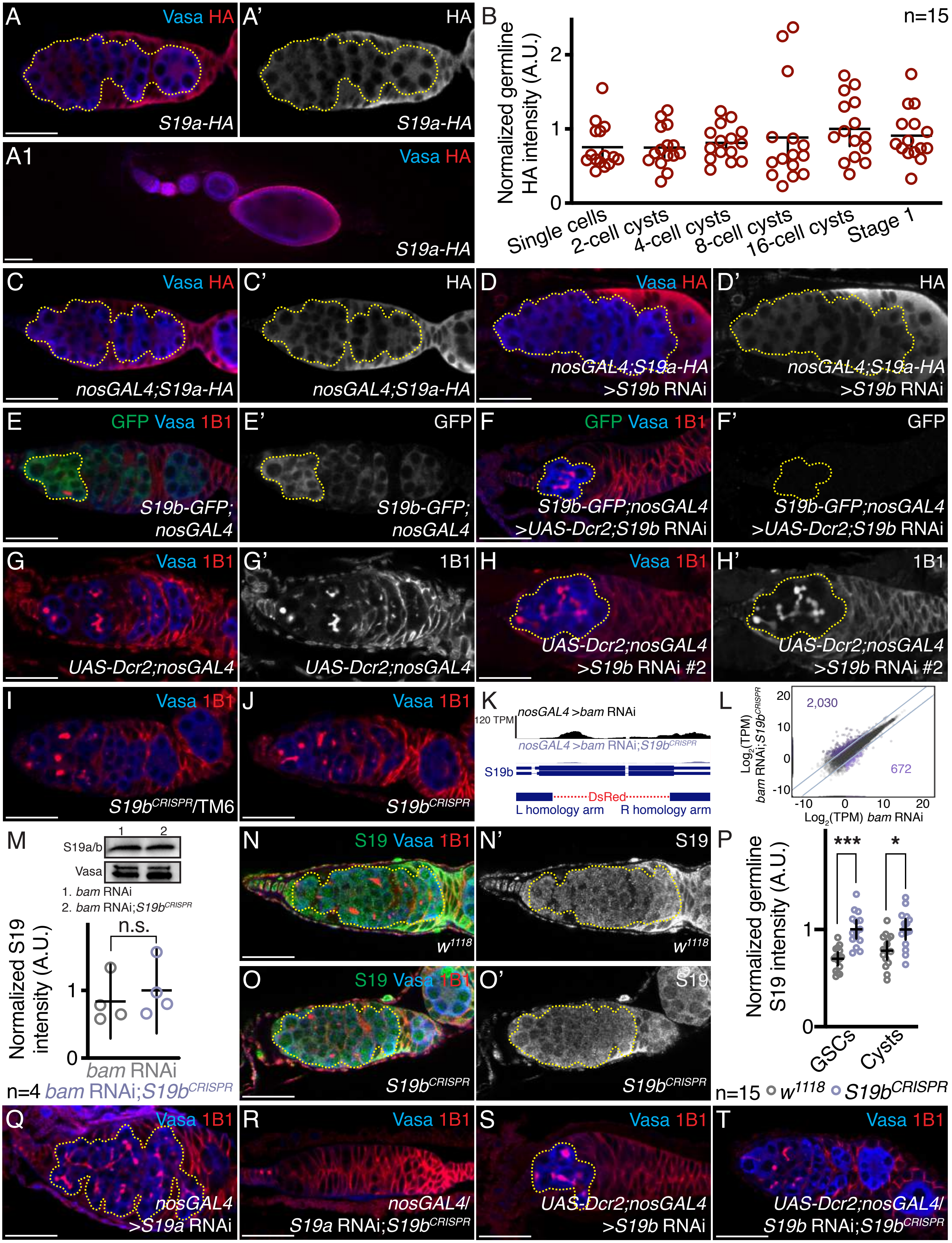

A Vasa HA S19a-HA
A' HA S19a-HA
A1 Vasa HA S19a-HA
B n=15
Normalized germline HA intensity (A.U.)
Single cells
2-cell cysts
4-cell cysts
8-cell cysts
16-cell cysts
Stage 1
C Vasa HA nosGAL4;S19a-HA
C' HA nosGAL4;S19a-HA
D Vasa HA nosGAL4;S19a-HA >S19b RNAi
D' HA nosGAL4;S19a-HA >S19b RNAi
E GFP Vasa 1B1 S19b-GFP; nosGAL4
E' GFP S19b-GFP; nosGAL4
F GFP Vasa 1B1 S19b-GFP;nosGAL4 >UAS-Dcr2;S19b RNAi
F' GFP S19b-GFP;nosGAL4 >UAS-Dcr2;S19b RNAi
G Vasa 1B1 UAS-Dcr2;nosGAL4
G' 1B1 UAS-Dcr2;nosGAL4
H Vasa 1B1 UAS-Dcr2;nosGAL4 >S19b RNAi #2
H' 1B1 UAS-Dcr2;nosGAL4 >S19b RNAi #2
I Vasa 1B1 S19bCRISPR/TM6
J Vasa 1B1 S19bCRISPR
K nosGAL4 >bam RNAi
120 TPM
nosGAL4 >bam RNAi;S19bCRISPR
S19b
DsRed
L homology arm
R homology arm
L Log2(TPM) bam RNAi;S19bCRISPR
Log2(TPM) bam RNAi
2,030
672
M S19a/b
Vasa
1. bam RNAi
2. bam RNAi;S19bCRISPR
n.s.
Normalized S19 intensity (A.U.)
bam RNAi
n=4 bam RNAi;S19bCRISPR
N S19 Vasa 1B1 w1118
N' S19 w1118
O S19 Vasa 1B1 S19bCRISPR
O' S19 S19bCRISPR
P Normalized germline S19 intensity (A.U.)
***
*
GSCs
Cysts
n=15 w1118 S19bCRISPR
Q Vasa 1B1 nosGAL4 >S19a RNAi
R Vasa 1B1 nosGAL4/ S19a RNAi;S19bCRISPR
S Vasa 1B1 UAS-Dcr2;nosGAL4 >S19b RNAi
T Vasa 1B1 UAS-Dcr2;nosGAL4/ S19b RNAi;S19bCRISPR

Figure 5-Supplement 2

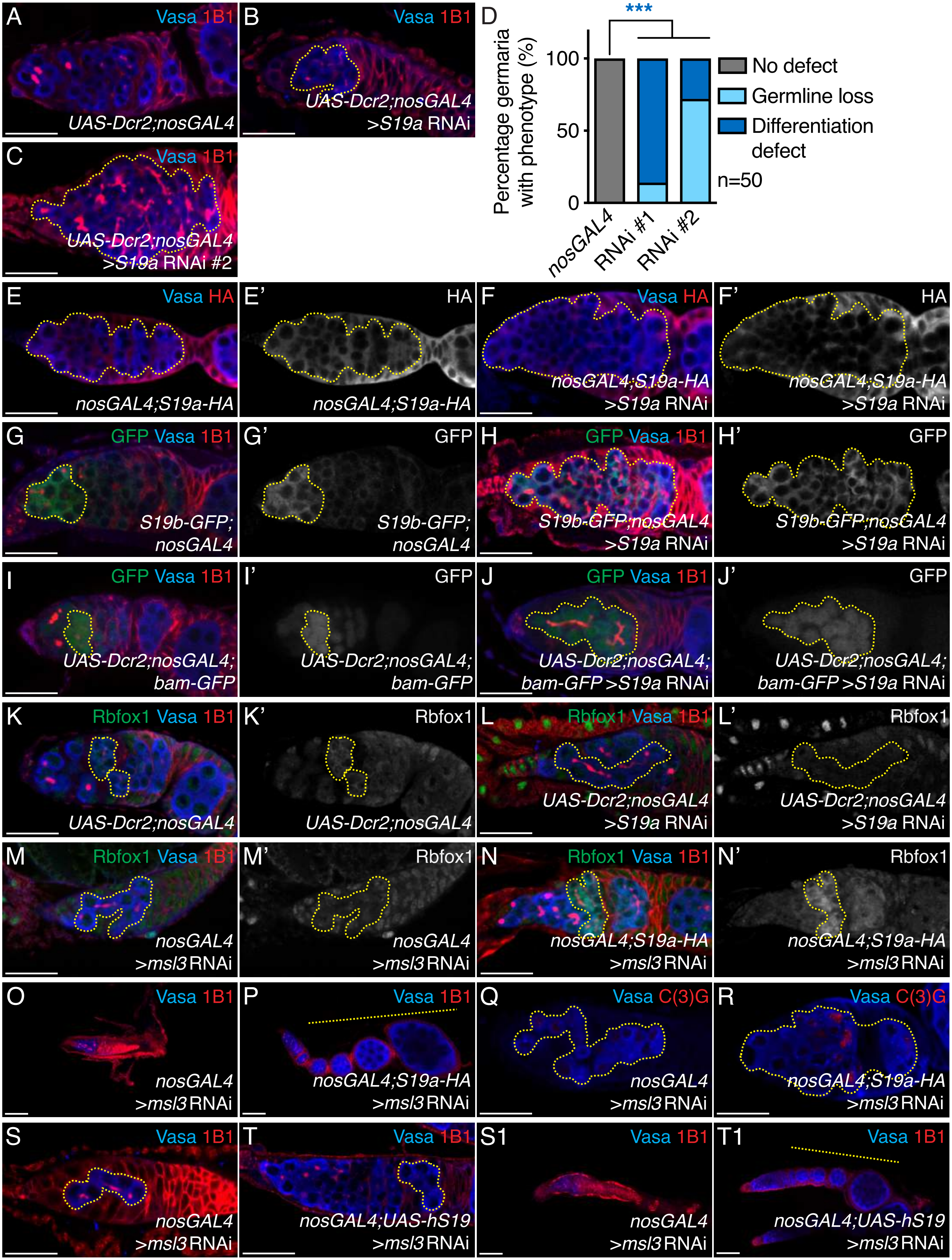

**Supplementary Figure Legends**

**Figure 1-Supplement 1. Set2 is required for proper cyst formation and Rbfox1 expression.**

(A-A') Control and (B-B') germline depleted *Set2* (line #2) germaria stained for Vasa (blue) and 1B1 (red) shows that *Set2* germline depletion results in irregular cysts (yellow dashed outline) (84% in *Set2* RNAi line #2 compared to 0% in *nosGAL4*; $p<2.2E-16$, n=50). Statistical analysis performed with Fisher's exact test. 1B1 channel is shown in A' and B'.

(C-C') Control and (D-D') germline depleted *Set2* germaria stained for H3K36me3 (green), Vasa (blue), and 1B1 (red) shows that *Set2* germline depletion results in decreased levels of H3K36me3 compared to control (yellow dashed outline and white arrow) ($0.1\pm0.1$ in *Set2* RNAi compared to $1.0\pm0.1$ in *nosGAL4*; $p<0.0001$, n=15). H3K36me3 channel is shown in C' and D'. Quantitation in (E), statistical analysis performed with Student t-test; *** indicates $p<0.001$.

(F-F') Control and (G-G') germline depleted *Set2* germaria stained for pMad (green), Vasa (blue), and 1B1 (red) shows that *Set2* germline depletion does not result in an increase in number of pMad positive germ cells compared to control (yellow dashed outline) ($1.1\pm1.1$ in *Set2* RNAi compared to $2.0\pm0.8$ in *nosGAL4*; $p=1.9E-5$, n=50). Statistical analysis performed with Student t-test. pMad channel is shown in F' and G'.

(H-H') Control and (I-I') germline depleted *Set2* germaria stained for Vasa (blue) and Bam (red) shows that *Set2* germline depletion results in an expansion of Bam positive germ cells compared to control (yellow dashed outline) (70% in *Set2* RNAi compared to 0% in *nosGAL4*; $p=4.1E-15$, n=50). Statistical analysis performed with Fisher's exact test. Bam channel is shown in H' and I'.

(J-J') Control and (K-K') germline depleted *Set2* germaria stained for Bruno (green), Vasa (blue), and 1B1 (red)shows that *Set2* germline depletion results in reduced levels of Bruno compared to control (yellow dashed outline and white arrow) ($0.5\pm0.1$ in *Set2* RNAi compared to $1.0\pm0.2$ in *nosGAL4*; $p=0.0024$, n=15). Bruno channel is shown in J' and K'. Quantitation in (L), statistical analysis performed with Student t-test; ** indicates $p<0.01$.

(M-M') Control and (N-N') germline depleted *Set2* germaria stained for Egl (green), Vasa (blue), and 1B1 (red) shows that *Set2* germline depletion results in aberrant Egl localization compared to control (yellow dashed outline) (90% in *Set2* RNAi compared to 4% in *nosGAL4*; $p=9E-16$, n=50) and improper oocyte specification (white arrow). Statistical analysis performed with Fisher's exact test. Egl channel is shown in M' and N'.

Scale bar for all images is 20 μm.

**Figure 2-Supplement 1. MSL3 is required in the germline and works with Set2**
(A) RNA-seq track showing that *msl3* is expressed during oogenesis. All tracks are set to scale to 7 TPM.

(B) Quantitation of frequency of germline MSL3-GFP expression in single cells, 2-cell cyst, 4-cell cyst, 8-cell cyst, 16-cell cyst, and stage 1 egg chambers (34% in single cells, n=50; 65% in 2-cell cyst, n=34; 55% in 4-cell cyst, n=33; 7% in 8-cell cyst, n=27; 4% in 16-cell cyst, n=26; and 0% in stage 1 egg chamber, n=20), showing that MSL3 is expressed during the mitotic and early meiotic stages of oogenesis.

(C-C') Heterozygous controls and (D-D') trans-allelic *msl3* mutant germaria stained for Vasa (blue) and 1B1 (red) shows that *msl3* mutants have irregular cysts (yellow dashed outline) (62% in $msl3^{1}/msl3^{MB}$ compared to 0% in $msl3^{1}$ and $msl3^{MB}$ heterozygotes; *p=1.7E-07*, n=72) and germline loss (38% in $msl3^{1}/msl3^{MB}$ compared to 0% in $msl3^{1}$ and $msl3^{MB}$ heterozygotes; *p<2.2E-16*, n=72). 1B1 channel is shown in C', and D'. Quantitation in Figure 2D.

(E-F) Heterozygous controls and (G) trans-heterozygous $Set2^{1}/+;msl3^{1}/+$ mutant germaria stained for Vasa (blue) and 1B1 (red) shows that trans-heterozygotes have severe germline loss compared to heterozygous control (100% in $Set2^{1}/+;msl3^{1}/+$ compared to 0% in $Set2^{1}$ heterozygotes and 0% in $msl3^{1}$ heterozygotes; *p<2.2E-16* for both, n=50). Statistical analysis performed with Fisher's exact test.

(H-H') Control and (I-I') germline depleted *Set2* germaria stained for GFP (green), Vasa (blue), and 1B1 (red) shows that *Set2* germline depletion results in aberrant MSL3-GFP localization compared to control (yellow dashed outline) (100% in *Set2* RNAi compared to 3.3% in *nosGAL4;msl3-GFP*; *p=5.2E-16*, n=30). Statistical analysis performed with Fisher's exact test. GFP channel is shown in H' and I'.

(J-J') Control and (K-K') germline depleted *msl3* germaria stained for H3K36me3 (green), Vasa (blue), and 1B1 (red) shows that *msl3* germline depletion results in unchanged levels of H3K36me3 compared to control (yellow dashed outline and white arrow) (0.9±0.1 in *msl3* RNAi compared to 1.0±0.1 in *nosGAL4*; *p=0.69*, n=15). H3K36me3 channel is shown in J' and K'. Quantitation in (L), statistical analysis performed with Student t-test; "n.s." indicates *p>0.5*.

Scale bar for all images is 20 µm.

**Figure 2-Supplement 2. MSL3 works independently of MSL complex in the ovaries**

(A-A') Control and (B-B') germline depleted *msl3* germaria stained for Vasa (blue) and 1B1 (red) shows that *msl3* germline depletion results in irregular cyst formation (yellow dashed outline) (87% in *msl3* RNAi compared to 0% in *nosGAL4*; *p<2.2E-16*, n=70) and germline loss (13% in *msl3* RNAi compared to 0% in *nosGAL4*; *p<2.2E-16*, n=70). 1B1 channel is shown in A' and B'. Quantitation in Figure 2D.

(C-C') Control and (D-D') germline depleted *msl3* germaria stained for pMad (green), Vasa (blue), and 1B1 (red) shows that *msl3* germline depletion does not result in an increase in number of pMad positive germ cells compared to control (yellow dashed outline) (1.0±0.9 in *msl3* RNAi compared to 2.0±0.7 in *nosGAL4*; *p<0.0001*, n=30). Statistical analysis performed with Student t-test. pMad channel is shown in C' and D'.

(E-E') Control and (F-F') germline depleted *msl3* germaria stained for Vasa (blue) and Bam (red) shows that *msl3* germline depletion results in an expansion of Bam positive germ cells compared to control (yellow dashed outline) (26% in *msl3* RNAi compared to 0% in *nosGAL4*; *p<3.5E-10*, n=50). Statistical analysis performed with Fisher's exact test. Bam channel is shown in E' and F'.

(G-G') Control and (H-H') germline depleted *msl3* germaria stained for Bruno (green), Vasa (blue), and 1B1 (red) shows that *msl3* germline depletion results in reduced levels of Bruno compared to control (yellow dashed outline and white arrow) (0.56±0.04 in *msl3* RNAi compared to 1.00±0.07 in *nosGAL4*; *p<0.0001*, n=15). Bruno channel is shown in G' and H'. Quantitation in (I), statistical analysis performed with Student t-test; ** indicates *p<0.01*.

(J-J') Control and (K-K') germline depleted *msl3* germaria stained for Egl (green), Vasa (blue), and 1B1 (red) shows that *msl3* germline depletion results in aberrant Egl localization compared to control (yellow dashed outline) (96% in *msl3* RNAi compared to 0% in *nosGAL4*; *p<2.2E-16*, n=50) and improper oocyte specification (white arrow). Statistical analysis performed with Fisher's exact test. Egl channel is shown in J' and K'.

(L) Percentage heterozygous controls and trans-allelic MSL complex mutants with no defect (gray), germline loss (light blue), and differentiation defect (dark blue). *msl1*, *msl2*, and *mle* mutants do not show irregular cyst formation or germline loss compared to respective heterozygous controls (100% in $msl1^{\gamma216}$/+;$msl1^{kmB}$/+ compared to 100% in $msl1^{\gamma216}$ and $msl1^{kmB}$ heterozygotes; *p=1*, n=50; 100% in $msl2^{227}$/+;$msl2^{kmA}$/+ compared to 100% and 100% in $msl2^{227}$ and $msl2^{kmA}$ heterozygotes; *p=1* for both, n=50; 100% in $mle^{1}$/+;$mle^{9}$/+ compared to 100% in $mle^{1}$ and $mle^{9}$ heterozygotes; *p=1* for both, n=50). Statistical analysis performed with Fisher's exact test on differentiation defect. No statistical significance was found.

Scale bar for all images is 20 µm.

**Figure 3-Supplement 1. ATAC members are required in the germline for proper cyst formation**

(A-A') Control, germline depleted (B-B') *NC2β*, (C-C') *Ada2a*, (D-D') *Atac1*, (E-E') *Atac2*, (F-F') *D12*, and (G-G') *wds* germaria stained for Vasa (blue) and 1B1 (red) shows that ATAC member germline depletion results in irregular cysts (yellow dashed outline) (24% in *NC2β* RNAi line #2, 90% in *Ada2a* RNAi, 25% in *Atac1* RNAi, 75% in *Atac2* RNAi, 77% in *D12* RNAi, and 68% in *wds* RNAi compared to 0% *nosGAL4*; $p<0.0001$ for all, n=50) and germline loss (21% in *NC2β* RNAi line #2, 10% in *Ada2a* RNAi, 20% in *Atac1* RNAi, 14% in *Atac2* RNAi, 3% in *D12* RNAi, and 22% in *wds* RNAi compared to 0% in *nosGAL4*; $p<0.05$ for *Atac1*, *Atac2*, and *wds* RNAi, $p>0.05$ for *Ada2a* and *D12* RNAi, n=50). 1B1 channel is shown in A', B', C', D', E', F', and G'. Quantitation in (H), statistical analysis performed with Fisher's exact test on differentiation defect; *** indicates $p<0.001$.

Scale bar for all images is 20 µm.

**Figure 3-Supplement 2. NC2β, an ATAC complex component, is required in the germline for proper differentiation and oocyte specification**

(A-A') Control and (B-B') germline depleted *NC2β* germaria stained for pMad (green), Vasa (blue), and 1B1 (red) shows that *NC2β* germline depletion does not result in an increase in number of pMad positive germ cells compared to control (yellow dashed outline) (1.4±1.3 in *NC2β* RNAi compared to 3.0±0.8 in *nosGAL4*; $p<0.0001$, n=50). Statistical analysis performed with Student t-test. pMad channel is shown in A' and B'.

(C-C') Control and (D-D') germline depleted *NC2β* germaria stained for Vasa (blue) and Bam (red) shows that *NC2β* germline depletion results in an expansion of Bam positive germ cells compared to control (yellow dashed outline) (70% in *NC2β* RNAi compared to 0% in *nosGAL4*; $p=4.1E-15$, n=50). Statistical analysis performed with Fisher's exact test. Bam channel is shown in C' and D'.

(E-E') Control and (F-F') germline depleted *NC2β* germaria stained for Bruno (green), Vasa (blue), and 1B1 (red) shows that *NC2β* germline depletion results in reduced levels of Bruno compared to control (yellow dashed outline and white arrow) (0.2±0.1 in *NC2β* RNAi compared to 1.0±0.3 in *nosGAL4*; $p<0.0001$, n=15). Bruno channel is shown in E' and F'. Quantitation in (G), statistical analysis performed with Student t-test; *** indicates $p<0.001$.

(H-H') Control and (I-I') germline depleted *NC2β* germaria stained for Egl (green), Vasa (blue), and 1B1 (red) shows that *NC2β* germline depletion results in aberrant Egl localization compared to control (yellow dashed outline) (72% in *NC2β* RNAi compared to 0% in *nosGAL4* germaria; *p=9.5E-16*, n=50) and improper oocyte specification (white arrow). Statistical analysis performed with Fisher's exact test. Egl channel is shown in H' and I'.

(J-J') Control and (K-K') germline depleted *NC2β* germaria stained for H3K36me3 (green), Vasa (blue), and 1B1 (red) shows that *NC2β* germline depletion results in unchanged levels of H3K36me3 compared to control (yellow dashed outline) ($1.1\pm0.5$ in *NC2β* RNAi compared to $1.0\pm0.5$ in *nosGAL4*; *p=0.65*, n=15). H3K36me3 channel is shown in J' and K'. Quantitation in (L), statistical analysis performed with Student t-test; "n.s." indicates *p>0.5*.

(M-N) Heterozygous controls and (O) trans-heterozygous $Atac2^{e03046}/+;msl3^{1}/+$ mutant germaria stained for Vasa (blue) and 1B1 (red) shows that trans-heterozygotes have severe germline loss compared to heterozygous controls (100% in $Atac2^{e03046}/+;msl3^{1}/+$ compared to 0% in $msl3^{1}$ and $Atac2^{e03046}$ heterozygotes; *p=1.6E-14* for both, n=25). Statistical analysis performed with Fisher's exact test on differentiation defect.

Scale bar for all images is 20 μm.

**Figure 4-Supplement 1. MSL3 regulates levels of meiosis-promoting genes and the germline enriched ribosomal protein, *RpS19b***

(A-A') Control and (B-B') germline depleted *msl3* germaria stained for RNA probes against *Rbfox1* (red) and DAPI (blue) shows that *msl3* germline depletion results in unchanged *Rbfox1* levels in the germline compared to control ($0.9\pm0.4$ in *msl3* RNAi compared to $1.0\pm0.2$ in *bam* RNAi;*hs-bam*; *p=0.005*, n=15). Statistical analysis performed with Student t-test. *Rbfox1* channel is shown in A' and B'.

(C) RNA-seq track showing that *sunn* is reduced upon germline depletion of *Set2*, *msl3*, and *NC2β*. All tracks are set to scale to 6 TPM.

(D) RNA-seq track showing that *ord* is reduced upon germline depletion of *Set2*, *msl3*, and *NC2β*. All tracks are set to scale to 5 TPM.

(E-E') Control and (F-F') germline depleted *msl3* germaria stained for GFP (green), Vasa (blue), and 1B1 (red) shows that *msl3* germline depletion results in lower and mislocalized GFP levels compared to control ($0.5\pm0.2$ in *msl3* RNAi compared to $1.0\pm0.2$ in *bam*

RNAi;*hs-bam*; *p<0.0001*, n=15). Statistical analysis performed with Student t-test. GFP channel is shown in E' and F'.

(G-G') Control and (H-H') germline depleted *msl3* germaria stained for RNA probes against *RpS19a* (red) and DAPI (blue) shows that *msl3* germline depletion results in unchanged *RpS19a* levels in the germline compared to control (1.16±0.12 in *msl3* RNAi compared to 1.00±0.12 in *bam* RNAi;*hs-bam*; *p=0.35*, n=15). Statistical analysis performed with Student t-test. *RpS19a* channel is shown in G' and H'.

(I-I') Control and (J-J') germline depleted *msl3* germaria stained for RNA probes against *RpS19b* (red) and DAPI (blue) shows that *msl3* germline depletion results in lower *RpS19b* levels in the germline compared to control (0.4±0.2 in *msl3* RNAi compared to 1.0±0.3 in *bam* RNAi;*hs-bam*; *p<0.0001*, n=15). Statistical analysis performed with Student t-test. *RpS19b* channel is shown in I' and J'.

Scale bar for all images is 20 μm.

**Figure 5-Supplement 1. RpS19a is in the germline and soma of *Drosophila* ovaries**
(A-A') *RpS19a-HA* germarium and (A1) ovariole stained for Vasa (blue) and HA (red) shows that HA expression is in soma and germline (0.8±0.1 in single cells, 0.8±0.1 in 2-cell cyst, 0.8±0.1 in 4-cell cyst, 0.9±0.2 in 8-cell cyst, 1.0±0.1 in 16-cell cyst, and 0.9±0.1 in stage 1 egg chamber; *p>0.5*, n=15). HA channel is shown in A'. Quantitation in (B), statistical analysis performed with one-way ANOVA; no statistical significance was found.

(C-C') Control and (D-D') germline depleted *RpS19b* germaria stained for Vasa (blue) and HA (red) shows that *RpS19b* germline depletion does not result in decreased RpS19a-HA compared to control (1.3±0.2 in *RpS19b* RNAi compared to 1.0±0.2 in *nosGAL4*; *p>0.05*, n=15). Statistical analysis performed with Student t-test. HA channel is shown in C' and D'.

(E-E') Control and (F-F') germline depleted *RpS19b* germaria stained for GFP (green), Vasa (blue) and 1B1 (red) shows that *RpS19b* germline depletion results in decreased RpS19b-GFP expression compared to control (0.6±0.3 in *RpS19b* RNAi compared to 1.0±0.2 in *nosGAL4*; *p=0.013*, respectively, n=15). Statistical analysis performed with Student t-test. GFP channel is shown in E' and F'.

(G-G') Control and (H-H') germline depleted *RpS19b* (line #2) germaria stained for Vasa (blue) and 1B1 (red) shows that *RpS19b* germline depletion results in accumulation of irregular cysts (yellow dashed outline) (32% in *RpS19b* RNAi line #2 compared to 0% in *nosGAL4*; *p=7.3E-6*, n=50). Statistical analysis performed with Fisher's exact test. 1B1 channel is shown in G' and H'.

(I) Heterozygous and (J) homozygous *RpS19b*$^{CRISPR}$ mutant germaria stained for Vasa (blue) and 1B1 (red) shows that *RpS19b* mutants do not show cyst defects (97% in *RpS19b* homozygotes compared to 100% in *RpS19b* heterozygotes; *p=1*, n=30). Statistical analysis performed with Fisher's exact test.

(K) Top: RNA-seq track showing that *RpS19b* reads are reduced in *bam* RNAi;*RpS19b*$^{CRISPR}$ compared to *bam* RNAi. All tracks are set to scale to 120 TPM. Below: Schematic of *RpS19b*$^{CRISPR}$ mutant design.

(L) Biplot of $Log_2$(TPM)*bam* RNAi;*S19b*$^{CRISPR}$ vs. $Log_2$(TPM)*bam* RNAi of *bam* RNAi. Light purple dots represent significantly downregulated transcripts and dark purple dots represent significantly upregulated transcripts in *bam* RNAi;*S19b*$^{CRISPR}$ ovaries compared with *bam* RNAi ovaries. Genes with four-fold or higher change were considered significant.

(M) Top: Western blot analysis of *bam* RNAi and *bam* RNAi;*S19b*$^{CRISPR}$ ovaries. The blot was stained for RpS19 and Vasa showing that RpS19 levels are not significantly decreased in *bam* RNAi;*RpS19b*$^{CRISPR}$ compared to *bam* RNAi (1.0±0.2 in *RpS19b*$^{CRISPR}$ compared to 0.8±0.2 in *bam* RNAi; *p=0.5594*, n=4. Bottom: Quantitation, statistical analysis performed with Student t-test; "n.s." indicates *p>0.5*.

(N-N') Control and (O-O') *RpS19b*$^{CRISPR}$ germaria stained for RpS19 (green), Vasa (blue), and 1B1 (red) shows that *RpS19b*$^{CRISPR}$ germaria do not have decreased RpS19 expression compared to control (1.0±0.2 in *RpS19b*$^{CRISPR}$ GSCs compared to 0.7±0.1 in $w^{1118}$ GSCs; *p<0.0001*, n=15; 1.0±0.2 in *RpS19b*$^{CRISPR}$ cysts compared to 0.8±0.2 in $w^{1118}$ cysts; *p=0.0023*, n=15). S19 channel is shown in N' and O'. Quantitation in (P), statistical analysis performed with Student t-test; * indicates *p.0.05* and *** indicates *p<0.001*.

(Q) Germline depleted *RpS19a* and (R) germline depleted of *RpS19a* in homozygous *RpS19b*$^{CRISPR}$ mutant germaria stained for Vasa (blue) and 1B1 (red) shows that *RpS19a* germline depletion in *RpS19b*$^{CRISPR}$ mutants results in germline loss compared to germaria with germline depletion of *RpS19a* (yellow dashed outline) (78% in *RpS19a* RNAi;*RpS19b*$^{CRISPR}$ compared to 100% in *RpS19a* RNAi; *p<2.2E-16*, n=50). Statistical analysis performed with Fisher's exact test.

(S) Germline depleted *RpS19b* and (T) germline depleted of *RpS19b* in homozygous *RpS19b*$^{CRISPR}$ mutant germaria stained for Vasa (blue) and 1B1 (red) shows that *RpS19b* germline depletion in *RpS19b*$^{CRISPR}$ mutants results in no defects compared to germaria with germline depletion of *RpS19b* (yellow dashed outline) (100% in *RpS19b* RNAi;*RpS19b*$^{CRISPR}$ compared to 20% in *RpS19b* RNAi; $p<2.2E-16$, n=50). Statistical analysis performed with Fisher's exact test.

Scale bar for all images is 20 µm.

**Figure 5-Supplement 2. RpS19a rescues *msl3* differentiation defect but not meiotic progression defect**

(A) Control and (B-C) germline depleted *RpS19a* (RNAi line #1 and 2) germaria stained for Vasa (blue) and 1B1 (red) shows that *RpS19a* germline depletion results in irregular cysts (yellow dashed outline) (90% in *RpS19a* RNAi line #1 and 60% in *RpS19a* RNAi line #2 compared to 0% in *nosGAL4*; $p<2.2E-16$ and $p=3.1E-15$, respectively, n=50) and germline loss (10% in *RpS19a* RNAi line #1 and 40% in *RpS19a* RNAi line #2 compared to 0% in *nosGAL4*; $p>0.05$ and $p=5.5E-9$, respectively, n=50). Quantitation in (D), statistical analysis performed with Fisher's exact test on differentiation defect; *** indicates $p<0.001$.

(E-E') Control and (F-F') germline depleted *RpS19a* germaria stained for Vasa (blue) and HA (red) shows that *RpS19a* germline depletion results in decreased RpS19a-HA expression compared to control (0.3±0.2 in *RpS19a* RNAi compared to 1.0±0.7 in *nosGAL4*; $p=0.005$ n=15). Statistical analysis performed with Student t-test. HA channel is shown in E' and F'.

(G-G') Control and (H-H') germline depleted *RpS19a* germaria stained for GFP (green), Vasa (blue) and 1B1 (red) shows that *RpS19a* germline depletion does not result in decreased RpS19b-GFP expression compared to control (1.7±0.5 in *RpS19a* RNAi compared to 1.0±0.3 in *nosGAL4*; $p<0.0001$, n=15). Statistical analysis performed with Student t-test. GFP channel is shown in G' and H'.

(I-I') Control and (J-J') germline depleted *RpS19a* germaria both carrying a *bam-GFP* transgene stained for GFP (green), Vasa (blue), and 1B1 (red) shows that *RpS19a* germline depletion results in irregular GFP positive cysts compared to control (yellow dashed outline) (80% in *RpS19b* RNAi compared to 0% in *nosGAL4*; $p=2.5E-13$, n=50). Statistical analysis performed with Fisher's exact test. GFP channel is shown in I' and J'.

(K-K') Control and (L-L') germline depleted *RpS19a* germaria stained for Rbfox1 (green), Vasa (blue), and 1B1 (red) shows that *RpS19a* germline depletion results in decreased levels of Rbfox1 in the germline compared to control (yellow dashed outline) (0.6±0.4 in

*RpS19a* RNAi compared to 1.0±0.6 in *nosGAL4*; *p=0.04*, n=15). Statistical analysis performed with Student t-test. Rbfox1 channel is shown in K' and L'.

(M-M') Control and (N-N') *RpS19a-HA* rescue germaria stained for Rbfox1 (green), Vasa (blue), and 1B1 (red) shows that addition of *RpS19a-HA* to *msl3* depletion ovaries results in increased Rbfox1 levels compared to control (1.0±0.8 in rescue compared to 0.2±0.04 in *msl3* RNAi; *p=0.0012*, n=15). Statistical analysis performed with Student t-test. Rbfox1 channel is shown in M' and N'.

(O) Control and (P) *RpS19a-HA* rescue germaria stained for Vasa (blue) and 1B1 (red) shows that addition of *RpS19a-HA* to *msl3* depletion ovaries results in an increased frequency of spectrosomes and cysts (86% in *RpS19a-HA* rescue compared to 18% in *msl3* RNAi; *p=1.5E-8*, n=50) and subsequent egg chambers compared to control (yellow dashed outline) (78% in *RpS19a-HA* rescue compared to 18% in *msl3* RNAi; *p=2.1E-9*, n=50). Statistical analysis performed with Fisher's exact test.

(Q) Control and (R) *RpS19a-HA* rescue germaria stained for Vasa (blue) and C(3)G (red) shows that rescue and control germaria both have aberrant C(3)G expression (yellow dashed outline and white arrows) (85% in *RpS19a-HA* rescue compared to 100% in *msl3* RNAi; *p=0.23*, n=50). Addition of *RpS19a-HA* does not rescue egg laying defects (28 eggs/female in *RpS19a-HA* compared to 0 eggs/female in *msl3* RNAi and rescue; *p<0.0001*, n=4). Statistical analysis performed with Fisher's exact test.

(S) Control and (T) *hRpS19-HA* rescue germaria stained for Vasa (blue) and 1B1 (red) shows that expression of *UAS-hRpS19-HA* in *msl3* germline depletion ovaries results in single cells and cysts compared to control (yellow dashed outline) (96% in *hRpS19-HA* rescue compared to 22% in *msl3* RNAi; *p=7.6E-9*, n=40). Statistical analysis performed with Fisher's exact test.

(S1) Control and (T1) *hRpS19-HA* rescue ovarioles stained for Vasa (blue) and 1B1 (red) shows that expression of *UAS-hRpS19-HA* in *msl3* germline depletion ovaries results in an increased frequency of subsequent egg chambers compared to control (yellow dashed outline) (100% in *hRpS19-HA* rescue compared to 15% in *msl3* RNAi; *p<2.2E-16*, n=40). Statistical analysis performed with Fisher's exact test.

Scale bar for all images is 20 μm.

**Supplementary methods**

RpS19b Homologous Arm 1

GGGTGTCGCCCTTCGCTGAAGCAGGTGGAATTCGTTTAGCCATGCCATCTGCGGA
GGGGTCTTATCGGCCAGTCGTATTTCCTGCGCCAGTTCATTGTCCAGCAGATAAGT
GGCCAAGTTGCGTCCAATGAAGCCGCAGCctgcgggaagaggaagagcagcgatgcagtcagcg
atgctgataagcggaagtgtggagtggggagcgggggcaggatgacgatgaggaggatgcagcgggaccacttacC
ACCCAAAATCAGAACAGTCGGCTTTTCAGACATGCTGCAGGCGCGACACGGTCAA
CCGGAATGCGAATGAGTCAAGTCGAAAGATATATATTTGTTAGCGCTCGAAATGCT
TTTTGTTGTTTTGTTTGGCTGTATTTCTAGTGTGACCatcccgtgatcattcttaaaacaagaaata
ccttcaatgctccacaagctggtcaccctatggcgctgtattcaaacctcttcgaatgctcttcaattagcgtagaacgttttatt
catttatacaaaccaaaatacagcagttctttgattcttactccatagggcttcagtgtcataaaattgtccttgttagtctacctta
aaccccctgtcaccaagaacagtagttttgctacattgcgttcctttccaagtttataggttaagttttcgaaaagaattgccaa
gttcaccaatactgcacgacaaaactatcggaaagtaatcctttggggtcaatgacctggtccctctttcaaacgtcacttgc
caatcgtccATTTTGACTAAAGTCAGCAAACTAACCGGAAGCAAACCTTTTTCTTCTCAA
AAGgtaagactttttaaatacatacatacatatatcggacccatgagccttaaaaacgtgctttcaagAACGTAGAG
AGCAACATGCCTGGAGTCACAGAATTCTTGCATGCTAGCGGCCGCGGACATAT

RpS19b Homologous Arm 2

TGCATAAGGCGCGCCTAGGCCTTCTGCAGCTCCATGATTATCACTACCTAAGATGA
TAATACCAATACCACGATGGATCATCTTTGAGTGCCTTGGGTTTTCAAAACATTCTC
AATAAATCGAAACCAAAAATAAAATTAAACAATTATATACATAATAAAATAACTTAATA
CTTGAAATATAAACGATTCGAAACTTTATTTTTGACACAAACACGCCGAATAGCAAG
AATCCAAGTTATAAAATGTTAGCTAGACCACAAATCTCCCGATTTTTGGTTGCCAGG
AGCTCTGAAACCGCGAACTATGACCTCGGCCTGCCACCAACCGCTGGTTTCATTTG
CTGCTTTCAAAACGTCATGCTCTTCGTTGGTCTGGCTAACGTCTTCCCCAGATTTAT
CCAGATTCCGCACGCCAAGATCGTGGTAAATATCCATCTCCTTGCAGCAGCGTAAT
TCATTCGGTGTGTTTTCACTGCAATGCAGTAGATGCAGGTGTATGTTCTTGCAGCT
GCGCGGAAGTTGGCTTAATATATGCTTTGGGATTAGATTAAAGGAGTCACAATGGG
ATCCTTGGGGTCGATCGGAGTCTGTAAGGAGCTCCATTTCTAGAAACACGTTGGTG
ACTTTTAGTTCGGTGGAGAATAGCCTGTACAGCTCCTCGCAGGTGAGCAGGCGGA
GCAGCGCCTGGTTGATGCGACCTTCGGGCAGAGCATGGTGCTCGCAATAGTGATC
TCTGGATATGGAATCATGTGGTATGTGCAGAGTATAGGCCTCCGAGGGCATGAAC
ATATTGTTGCCAAAAAGAAATCGCAGGCTCTTCACTTCAGAGCTCCTGAAGGCTTG
TCGGATGATctgtaggagagaactggcaagtaaagaccaaaagaagtcctggatgtaaggggaacctcacCT
CTCTCATGTCACTAGCTCGAGGCTCTTCCGTCAATCGAGTTCAAG
